# Identification of MARVELlous Protein Markers for *Phytophthora infestans* Extracellular Vesicles

**DOI:** 10.1101/2025.04.11.648357

**Authors:** Susan Breen, Hazel McLellan, Wei Wang, Shumei Wang, Lydia Welsh, Jasmine Pham, Stephen C Whisson, Petra C Boevink, Paul RJ Birch

## Abstract

Extracellular vesicles (EVs) are released from cells by unconventional secretion, but little is known about the biogenesis routes, composition or cargoes of EVs from fungal or oomycete plant pathogens. We investigated the proteome of EV-associated proteins secreted by the oomycete *Phytophthora infestans*, cause of potato late blight disease. We found that vesicle-associated proteins, transmembrane proteins and RxLR effectors, which are delivered into host cells to suppress immunity, were enriched in the EV proteome. By contrast, the EV-independent secreted proteome was enriched in cell wall modifying enzymes and apoplastic effectors which act outside plant cells. Two proteins each containing two tetraspanning MARVEL domains, PiMDP1 and PiMDP2, were associated with *P. infestans* EVs. PiMDP1 and PiMDP2 were co-buoyant with RxLR effectors in sucrose density fractions containing EVs and co-localised frequently with each other and with RxLRs at vesicles within pathogen hyphae grown *in vitro* and during infection. Interestingly, PiMDP2, which is up-regulated during the early biotrophic phase of infection, accumulates at the haustorial interface, a major site of effector secretion during infection. We argue that PiMDP1 and PiMDP2 are molecular markers that will facilitate studies of the biogenesis and secretion of infection-associated *P. infestans* EVs.

## 1. INTRODUCTION

*Phytophthora infestans* is an economically important oomycete plant pathogen. It is the causal agent of late blight disease of potato and tomato and was responsible for the Irish potato famines in the mid-1800s (Kamoun et al. 2015). It is still relevant today, causing devastating crop loses and requiring the use of expensive and environmentally unfriendly chemical sprays (Tiwari et al. 2021). To protect food security, it is crucial to understand the mechanisms by which *P. infestans* causes disease.

An important infection strategy of *P. infestans* is its ability to produce virulence factors called effectors which are usually small proteins secreted by the pathogen. These can be apoplastic effectors, which are conventionally secreted and act in the extracellular space to inhibit host enzymes, degrade substrates or mask the presence of pathogen associated molecular patterns (PAMPs) (Wang et al. 2017; 2018; Sabbadin et al. 2021); or they can be cytoplasmic effectors which act inside host cells to manipulate and interfere with plant defences (Wang et al. 2023c). The most important class of cytoplasmic effectors in *P. infestans* contain the conserved motif Arg-any amino acid-Leu-Arg (RxLR), which are encoded by more than 500 genes in the *P. infestans* genome (Haas et al. 2009). The RxLR motif is required for effector translocation into living plant cells (Whisson et al. 2007). RxLR effectors have been reported to be non-conventionally secreted from pathogen haustoria (Wang et al. 2017; 2018). Cytoplasmic effectors from the fungal pathogen *Magnaporthe oryzae* are also exported by an unconventional secretion pathway (Giraldo and Valent 2013). Furthermore, cytoplasmic effectors from both *M. oryzae* and *P. infestans* have both been demonstrated to enter plant cells via Clathrin-mediated endocytosis after being observed in vesicle-like compartments in the pathogens’ specialised secretory structures (Oliveira-Garcia et al. 2023; Wang et al. 2023b).

Extracellular vesicles (EVs) are small lipid bilayer nanoparticles naturally released by cells in a form of non-conventional secretion. EVs have been extensively studied in mammalian systems and exist in several different forms; including exosomes released during the fusion of multivesicular bodies with the plasma membrane (PM); microvesicles produced directly from the PM; and autophagosomes released from cells undergoing apoptosis (Cai et al. 2021). EVs are involved in cell-to-cell communication within organisms but are also implicated in interactions between organisms and could play an important role during plant pathogen interactions, as demonstrated recently in the interaction between model plant *Arabidopsis* and fungal pathogen *Botrytis cinerea* (He et al. 2023; Wang et al. 2024).

To study EVs, it is crucial to identify marker proteins to enable the purification and identification of sub-populations of EVs; as well as to allow investigation of biogenesis pathways. In mammalian and plant systems tetraspanin proteins, which have four transmembrane (TM) domains and are integral membrane proteins, have been extensively used for this purpose. Plants have several different subtypes of EVs, including those associated with the markers TET8 or PEN1, which can be differentially purified using these membrane proteins (Cai et al. 2018; Rutter and Innes 2017; Cai et al. 2021). However, tetraspanin orthologues have not been identified in all EVs, including those from many fungi, so other membrane marker proteins such as SUR7 have been proposed (Dawson et al. 2020). SUR7 is a four TM containing protein normally located to MCC1/eisosomes in fungi. Eisosomes are stable membrane invaginations which serve as specific membrane microdomains thought to be enriched in membrane transport and endocytosis (Lanze et al. 2020). SUR7 was found to be enriched in EVs from *Candida albicans* and the fungal pathogen of plants, *Zymoseptoria tritici* (Dawson et al. 2020; Hill and Solomon 2020).

Little is known about EVs from oomycete plant pathogens. One study has indicated that tetraspanins produced by oomycetes may act as PAMPs in plant hosts (Zhu et al. 2023) and another showed that applying EVs to leaf tissue during infection can promote pathogenicity (Fang et al. 2021). Here, ultracentrifugation-mediated EV purification and liquid chromatography-mass spectrometry were used to determine an EV-associated proteome resulting in the identification of MARVEL domain-containing TM proteins as candidate EV membrane marker proteins. The MARVEL-containing markers are co-buoyant in sucrose density fractions with RxLR effectors and co-localise with them at vesicles in the pathogen both *in vitro* and during infection. These findings can be subsequently used by the community to investigate the roles of EVs in pathogenicity in *P. infestans*.

## 2. MATERIALS AND METHODS

### 2.1 *Phytophthora infestans* growth, plant growth and plant inoculation

*Phytophthora infestans* strain 3928A and transgenic lines made from this strain were cultured on Rye Agar (Welsh and Whisson 2025) supplemented with Ampicillin (100 µg/ml) and Pimaricin (0.001 %). Geneticin (10 µg/ml) was added to select for and maintain transgenic lines. *Nicotiana benthamiana* and *Solanum tuberosum* susceptible cultivar Maris Piper were grown in glasshouses in a 16/8-hour light-dark cycle. Plant leaves were inoculated on the abaxial surface with 10 µl drops of sporangia solution obtained by flooding a plate of 14 day old culture with sterile distilled water (SDW), scraping with a glass spreader and filtering through a 70 µm cell strainer into a 50 ml falcon tube. Sporangia solutions were centrifuged at 2700 rpm for 10 minutes and pellets were resuspended in fresh SDW, counted on a cell counter and adjusted to a concentration of 1000 sporangia per 10 µl drops.

### 2.2 Endo-vesicle Isolation from *P. infestans*

Endo-vesicles were isolated using 1–2 g of mycelia homogenized in an equal volume of isolation buffer (150 mM Na-HEPES, pH 7.5, 10 mM EDTA, 10 mM EGTA, 17.5% [w/w] sucrose, 7.5 mM KCl, 10 mM DTT, 1× PIC-W, 1× PIC-D, and 1× E-64) (Reynolds et al. 2014; Wang et al. 2023b). Cellular debris was removed by centrifugation at 2,000 × g for 20 minutes at 4 °C. The resulting supernatant was transferred into pre-chilled 1.5 mL Eppendorf tubes. The supernatant was divided into two equal volumes. One portion was treated with Triton X-100 to a final concentration of 1%, while the other received an equal volume of isolation buffer. Both samples were incubated on ice for 30 minutes with gentle shaking twice during incubation. Following this, they were ultracentrifuged at 100,000 × g for 60 minutes. The resulting pellets were resuspended in 2× SDS loading buffer for subsequent immunoblotting.

### 2.3 EV Isolation

14-day old Rye agar plates were flooded with 20 ml of amended lima bean (ALB) liquid media supplemented with the appropriate antibiotics scraped with a glass spreader into a sterile culture bottle through a 70 µm cell strainer and grown for 3 days in the dark without shaking. Mycelia were transferred into fresh culture bottles containing 15 ml minimal media and 15 ml ALB which had been pre-cleared by ultracentrifugation at 100 000 × g for 2.5 hours. Cultures were grown for a further 2 days. The culture filtrate (CF) was obtained by removing the mycelia with a 70 µm cell strainer. The CF was centrifuged for 10 minutes at 2000 × g, filtered through a 0.45 μm syringe filter, followed by a further spin at 10,000 × g for 30 minutes to remove all cellular debris. The CF was then ultracentrifuged at 100,000 × g for 1 hour at 4 °C (Beckman Coulter Type50.2 Ti rotor) to give the supernatant (100S) and crude pellet (100P) samples.

### 2.4 Transmission Electron Microscopy (TEM)

A 25 μl aliquot of EV 100P sample was mixed with 25 μl of 4% paraformaldehyde (PFA) in 0.1 M sodium phosphate pH 7.4 to give a final concentration of 2% PFA. 25 μl of this was pipetted onto a piece of parafilm and a carbon-coated, formvar, 300 mesh, nickel grid was placed on top and incubated at room temperature for 30 minutes. Grids were washed with 1 X PBS twice then incubated for 10 minutes with 50 μl 1% glutaraldehyde. Grids were washed with 100 μl sterile water seven times and stained with 4% uranyl oxalate for 10 minutes at room temperature. Grids were transferred onto two 50 μl drops of uranyl acetate (0.15% methylcellulose and 0.4% uranyl acetate) sequentially and left on the second drop for 10 minutes on ice. Excess liquid was wicked off using filter paper and grids were inspected and imaged with a JEOL 1400, at 80 kV.

### 2.5 Nanoparticle Tracking (NTA)

Particle size and concentration of EV pellet fractions was determined using a NanoSight light microscope NT10 with a 638 nm red laser or a NanoSite NS300 with a 488nm blue laser (Malvern Instruments). EV pellet samples were diluted between 1:4 to 1:6 with 1x PBS, filter sterilised using a 0.22 μM syringe filter (Millipore). NanoSight NTA 2.3 or NanoSite NTA3.4 software was used with settings of camera level 7-10, detection threshold 8-11, at room temperature with 3 to 5 replicate 30 second or 1-minute reads to collect data.

### 2.6 FM4-64 staining

Pellet samples were mixed 1:1 with FM4-64 for a final dye concentration of 20 µg/ml and imaged on microscope slides with the 561nm laser on a Nikon A1R confocal microscope using a x60 Nikon CFI Plan Apochromat VC WI water immersion lens with an NA of 1.2. Emissions were collected at 570-620 nm and the pinhole was set to 1 airy unit NIS-Elements or ImageJ were used to process the images.

### 2.7 Proteomics Sample Processing

Samples were four independent biological replicates of each of the *P. infestans* culture filtrate EV pellet and the supernatant from the 100,000 × g ultracentrifugation spin. Four independent biological replicates of pellets and 100,000 × g supernatants from media-only control spins, not containing *P. infestans*, were also used and treated in the same way. For the supernatant samples 2 ml of each was subjected to methanol-chloroform precipitation and pellets were air dried. All samples were resuspended in 50 μl 0.25% Rapigest^TM^ SF (Waters) in 50 mM ammonium bicarbonate. Protein concentration was determined by taking 2.5 μl of each sample and carrying out a scaled down version of the BCA assay (Pierce) according to manufacturer’s instructions. DTT was added to the remaining sample to a final concentration of 5 mM and samples were boiled at 100 °C for 5 minutes. Samples were passed to the ‘FingerPrints’ Proteomics Facility, School of Life Sciences, University of Dundee where they were subsequently processed. Samples were alkylated by adding 1.5 µl (500 mM Iodoacetamide) and incubated for 30 min the dark at RT. Digestion was achieved by adding 56 µl of ammonium bicarbonate containing 7.5 µg of trypsin/sample and incubation at 37 °C for 14 hrs. Peptides were acidified with TFA (0.5% final concentration) and incubated at 37 °C for 60 min to precipitate Rapigest. Samples were finally centrifuged at 13,000 rpm for 10 min, and supernatant containing peptides were transferred to new vials. Samples were resuspended in 50 µl 1 % formic acid and stored at -22 °C prior to LC-MS analysis.

### 2.8 LC-MS/MS

Analysis of peptide readout was performed on a Q Exactive™ plus, Mass Spectrometer (Thermo Scientific) coupled to a Dionex Ultimate 3000 RS (Thermo Scientific). LC buffers used were the following: buffer A (0.1 % formic acid in Milli-Q water (v/v)) and buffer B (80 % acetonitrile and 0.1 % formic acid in Milli-Q water (v/v). An equivalent of 1.25 µg of peptides from each sample were loaded at 10 μl/min onto a trap column (100 μm × 2 cm, PepMap nanoViper C18 column, 5 μm, 100 Å, Thermo Scientific) equilibrated in 0.1 % TFA for 15min. The trapping column was washed for 6 min at the same flow rate with 0.1 % TFA and then switched in-line with a Pharma Fluidics, 200 cm, µPAC nanoLC C18 column, equilibrated with buffer A at a flow rate of 300 nl/min for 30 min. The peptides were eluted from the column at a constant flow rate of 300 nl/min with a linear gradient from 1 % buffer B to 3.8 % buffer B in 6 min, from 3.8 % B to 12.5 % buffer B in 40 min, from 12.5 % buffer B to 41.3 % buffer B within 176 min and then from 41.3 % buffer B to 61.3 % buffer B in 34 min. The gradient was finally increased from 61.3 % buffer B to 100 % buffer B in 10 min, and the column was washed at 100 % buffer B for 10 min. Two blanks were run between each sample to reduce carry-over. The column was kept at a constant temperature of 50 °C. Q-exactive plus was operated in positive ionization mode using an easy spray source. The source voltage was set to 2.2 Kv and the capillary temperature was 275 °C. Data were acquired in Data Independent Acquisition (DIA) Mode as previously described (Doellinger et al. 2020), with little modification. A scan cycle comprised a full MS scan (m/z range from 345-1155), resolution was set to 70,000, AGC target 3 × 10^6^, maximum injection time 200 ms. MS survey scans were followed by DIA scans of dynamic window widths with an overlap of 0.5 Th. DIA spectra were recorded at a resolution of 17,500 at 200 m/z using an automatic gain control target of 3 × 10^6^, a maximum injection time of 55 ms and a first fixed mass of 200 m/z. Normalised collision energy was set to 25 % with a default charge state set at 3. Data for both MS scan and MS/MS DIA scan events were acquired in profile mode.

### 2.9 Proteomics Data Analysis

Analysis of the DIA data was carried out using Spectronaut (version 17.4.230317.55965, Biognosys, AG). The directDIA workflow, using the default settings (BGS Factory Settings) with the following modifications was followed: decoy generation set to mutated; Protein LFQ Method was set to QUANT 2.0 (SN Standard) and Precursor Filtering set to Identified (Qvalue); Cross-Run Normalization was unchecked; Precursor Qvalue Cutoff and Protein Qvalue Cutoff (experimental) were set to 0.01; Precursor PEP Cutoff set to 0.1 and Protein PEP Cutoff set to 0.75; Protein Qvalue Cutoff (Run) set to 0.05. For the Pulsar search, the settings were: maximum 2 missed trypsin cleavages; PSM, Protein and Peptide FDR levels set to 0.01; scanning range set to 300-1800 m/z and Relative Intensity (Minimum) set to 5%; cysteine carbamidomethylation set as fixed modification and acetyl (N-term), deamidation (asparagine, glutamine), dioxidiation (methionine, tryptophan), glutamine to pyro-Glu and oxidation of methionine set as variable modifications. Separate searches were made against the fasta files for *Phytophthora infestans* (strain T30-4; UP000006643, 17,610 entries) and *Phaseolus vulgaris* (UP000000226; 30,501 entries). Analysis of differential protein abundance for the *P. infestans* data was carried out in Excel with the Spectronaut output. Peptides that appeared to represent *P. infestans* proteins were removed if they were also identified in greater than 3 out of the 8 media control samples as an additional control for misidentification. Positive protein identification was confirmed only if there were >1 peptide identified over 3 out of 4 biological replicates. Perseus version 2.0.6.0 (http://www.coxdocs.org/doku.php?id=perseus:start) was subsequently used to do PCA analysis and produce volcano plot (2-sided t-test with 250 randomisations with a permutation based FDR calculation, FDR= 0.05, S0= 0.1). If proteins were not detected in either P or S samples they were given a value of zero in those samples to include them in the volcano plot.

### 2.10 Construction of *P. infestans* transgenic lines

The vector pmCitrine:mCherry used to co-express proteins of interest tagged with mCitrine and mCherry each under the control of the *Bremia lactucae* Ham34 promotor (Ham34P) and terminator (Ham34T) was based on pMCherryN (Ah-Fong and Judelson 2011) with the mCitrine cassette cloned into the Sac I site. Coding sequences of Pi04314 (XM_002905913.1), Pi09216 (XM_002903466.1), Pi13660 (XM_002899798.1), Pi13661 (XM_002899799.1), PiPE1 (XM_002909508.1), PiGNB2L (XM_002903143.1), PiRab7 (XM_002906189.1), PiCoronin (XM_002908188.1) and PiAVRblb1 (XM_002895005.1), were amplified from *P. infestans* isolate 3928A cDNA with the primers shown in Supplementary Table S1 and were cloned as either C or N terminal protein fusions to the fluorescent proteins using restriction enzyme cloning and standard molecular biology techniques. Plasmids were sequenced to confirm the correct inserts and Plasmid prepped using the Promega PureYield™ Plasmid Maxiprep System. Transformation of *P. infestans* isolate 3928A was performed using a modified polyethylene glycol-CaCl_2_-lipofectin protocol (Judelson et al. 1991; Welsh and Whisson 2025). Previously published transformants were also used for PiEPIC1 (Wang et al. 2017), PiManl (Ah-Fong and Judelson 2011), PiAvr3a (Whisson et al. 2007) and PiSCR74 (Lin et al. 2020).

### 2.11 Triton X-100 Treatments

Prior to the 100, 000 xg ultracentrifugation the CF was split into two equal volumes, one was treated with Triton X-100 detergent at a final concentration of 1%, while the other had the same volume of SDW added. The tubes were mixed and incubated on ice for one hour before ultracentrifuging at 100, 000 × g for one hour (Beckman Coulter, Type50.2 Ti rotor). Pellets were resuspended in equal volumes of 2 ×SDS protein loading buffer and equal volumes were loaded on SDS PAGE gels.

### 2.12 Sucrose Density Gradients

The crude pellet from the 100,000 × g ultracentrifugation was resuspended into 1 ml 1x PBS and adjusted to make a 10% sucrose solution which was top loaded onto a sucrose gradient composed of 10, 20, 30, 40, 50, 60 and 70% sucrose solutions of 1.6 ml each, made up with 1x PBS plus Protease inhibitor tablet. Samples were ultracentrifuged for 16 hours at 4 °C at 100,000 xg with no brake (Beckman Coulter, SW41 rotor). Six fractions of 1.8 ml each were taken, the percentage of sucrose measured using a portable refractometer (MRC, REF108 0-80 % Brix, 1 %) and used to calculate sucrose density. The tubes containing the fractions were topped up with 1 x PBS and then ultracentrifuged at 100,000 × g for 1 hour at 4 °C (Beckman Coulter, Type50.2 Ti rotor). The resulting pellets were resuspended in either 40 μl of 1 × PBS for TEM or 2xSDS PAGE loading buffer for Western blotting.

### 2.13 Western blotting

Samples were run on 10% SDS PAGE gels to separate samples and blotted onto Nitrocellulose membranes. Membranes were blocked in 4 % milk in 1 x PBST (0.05 % Tween 20) for one hour at room temperature. Primary antibodies for RFP/ mCherry (Chromotek, 58F) and GFP/ mCitrine (Roche, 11814460001) were incubated overnight at 4 °C in blocking buffer at a dilution of 1:4000 or 1:2000 respectively. Membranes were washed five times in 1 × PBST and the secondary antibodies IRDye® 680RD Goat anti-Rat IgG, IRDye® 800CW Goat anti-Rat IgG, IRDye® 800CW Goat anti-Mouse IgG or IRDye® 680RD Goat anti-Mouse IgG (LICORbio) were added. The membranes were incubated for 1 hr at room temperature in the dark in blocking buffer at a dilution of 1:5000. Blots were developed on a LiCOR ODYSSEY® DLx and quantified using LICOR Image Studio™ Software.

### 2.14 Phylogenetic analysis

Candidate MDP orthologs were identified by reciprocal best blast match (RBBM). Multiple sequence alignment was undertaken in Clustal omega and phylogenetic trees were generated using Phylogeny.fr (http://www.phylogeny.fr/) (Dereeper et al. 2008). Gene orientation and spacing was recorded in the genomes where both candidate MDP1 and MDP2 orthologues were identified. AlphaFold2 was used to predict the structures of PiMDP1 and PiMDP2 (https://colab.research.google.com/github/sokrypton/ColabFold/blob/main/AlphaFold2.ipynb).

### 2.15 qRT-PCR

Gene expression of PiMDP1, PiMDP2 and the RxLR Pi04314 was monitored using an infection time-course of *P. infestans* isolate 3928A on Maris Piper potato leaves. RNA was extracted using a RNeasy Plant mini kit (Qiagen). Potential DNA contamination was removed using the Turbo DNA-free kit (Thermo Fisher) and cDNA was synthesised using a Superscript III kit (Invitrogen). qRT-PCR was carried out using Power SYBR™ Green PCR Master Mix (Thermo Fisher) and the primers shown in Supplementary Table S1, on a StepOne Real-Time PCR System (Applied Biosystems). Expression was normalised to the geometric mean of 3 housekeeping genes: ACTIN (PITG_15117; EEY63399), Casein kinase (PITG_02745; EEY64204), Kelch domain repeat (PITG_09862; XM_002902372.1) using the 2^-ΔΔCt^ method (Lacomme et al. 2003).

### 2.16 Confocal Microscopy

*P. infestans* hyphae grown in ultracentrifuged ALB media on microscope slides were imaged at 2-3 dpi and leaf infections on either potato or *N. benthamiana* leaves were imaged at 3-5 dpi. A Nikon A1R confocal microscope was used. Samples on slides were imaged with a x60 Nikon CFI Plan Apochromat VC water immersion lens (NA 1.2) while infections were imaged with a x40 NIR Apo DIC water dipping lens (NA 0.8). The 514 nm line from an argon laser was used to excite the mCitrine with emissions collected at 520-550 nm. A 561nm laser was used to excite the mCherry and emissions collected at 570-620 nm. The pinhole was set to 1 airy unit for the 561 nm line. NIS-Elements or ImageJ were used to process and quantify the images. Fluorescence intensity plots drawn to show (co-)localisation in vesicle-like bodies were taken from single optical sections. HM /PM ratio measurements were taken by measuring the mean fluorescence intensity of comparably-sized regions of interest drawn to encompass the PM or HM in single optical sections from the same z series. Counts to show (co-)localisation of visually detectable fluorophores in vesicles from *in vitro* grown hyphae were performed manually from a minimum of 12 confocal images for each transformant and expressed as a percentage of the total number of vesicles counted.

## 3. RESULTS

### 3.1 *P. infestans* secretes EVs when grown *in vitro*

To determine whether *P. infestans* produces EVs we aimed to isolate EVs from *in vitro* cultures of wildtype *P. infestans* strain 3928A using ultracentrifugation (UC) methods (Figure 1a) broadly similar to those used for other filamentous plant and animal pathogens (Hill and Solomon 2020; Cai et al. 2021). Following UC, our crude pellets were resuspended and examined by Nanoparticle tracking analysis (NTA) and transmission electron microscopy (TEM) to verify the presence of EVs. NTA analysis showed that each of the pellets recovered by UC of *P. infestans* culture media contained an average equivalent particle concentration of 49.13 E8 particles/ml, with an average mean diameter of 202 nm and an average mode diameter of 100 nm, whereas the pellets derived from media without *P. infestans* lacked particles of a similar size or concentration but contained a low level of background particulate matter from the growth media (Figure 1b). TEM negative stain imaging showed the presence of typical cup-shaped vesicle structures of various sizes in the *P. infestans* culture media crude pellets (Figure 1c, Supplementary Figure S1a-b), whereas no vesicles were observed in the pellets of media-only samples generated using the same UC method (Supplementary Figure S1c). Staining of the *P. infestans* culture media crude pellet samples with lipophilic dye FM4-64 indicated that theys contain membrane-bound nanoparticles (Supplementary Figure S1d), whereas the uninoculated media-only pellets showed no FM4-64 staining (Supplementary Figure S1e).

**Figure 1:**
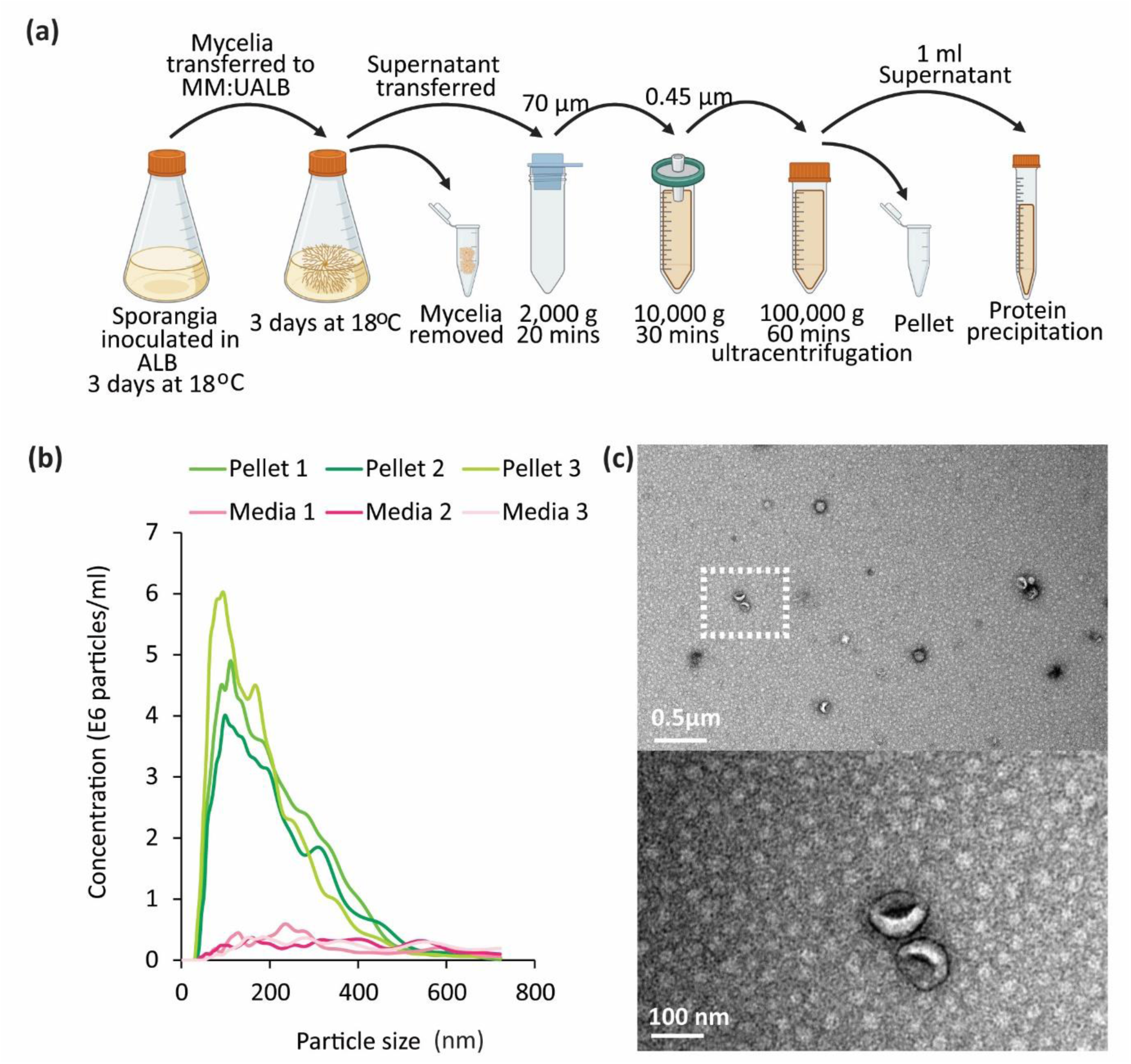
*Phytophthora infestans* secretes extracellular vesicle like particles into growth media. (a) Schematic representation of the ultracentrifugation protocol used for nanoparticle isolation to generate the crude pellet from *P. infestans* inoculated media. Schematic made in BioRender. (b) Nanoparticle tracking (NTA) data showing the size distribution of particles in the crude nanoparticle pellet from *P. infestans*-inoculated media and the pellet from uninoculated control media. Results from three independent replicates of each inoculated and uninoculated control are shown in the line graph. (c) Transmission electron microscopy negative stain images showing vesicles present within the nanoparticle crude pellet isolated from *P. infestans* inoculated media. Top image scale bar represents 0.5 µm. Dashed box represents the location of the higher magnification image below where the scale bar represents 100 nm.

### 3.2 Proteomics identifies an enrichment of EV-associated proteins

Confident that we were indeed obtaining EVs isolated from *P. infestans,* we proceeded to analyse the proteome of four independent replicates of the crude EV pellets and compared them to the proteins precipitated from the supernatants after the EVs had been removed by ultracentrifugation. The samples were all treated with Rapigest detergent to maximise recovery of integral membrane proteins possessing transmembrane (TM) domains. This approach yielded a starting point of 1908 proteins (Supplementary Table S2a). Low confidence proteins only represented by a single peptide and *P. infestans* proteins also present in more than 3 out of 8 of the media-only control samples were removed. This left 1326 high confidence proteins present in at least 3 out of the 4 biological replicates (Supplementary Table S2b). The EV pellet contained 450 unique proteins while the supernatant had only 23 unique proteins and there was an overlap of an additional 853 proteins (Figure 2a). PCA multiple component analysis showed that the pellet (EV-associated) and supernatant (EV-independent) samples clustered in their respective groups (Supplementary Figure S2a). GO term analysis of the proteins identified in the pellet samples show enrichments of proteins belonging to the functional categories of Metabolic enzymes, RNA metabolism and Translation, as well as Membrane traffic, Protein modifying enzymes and Transporters (Supplementary Figure S2b).

**Figure 2:**
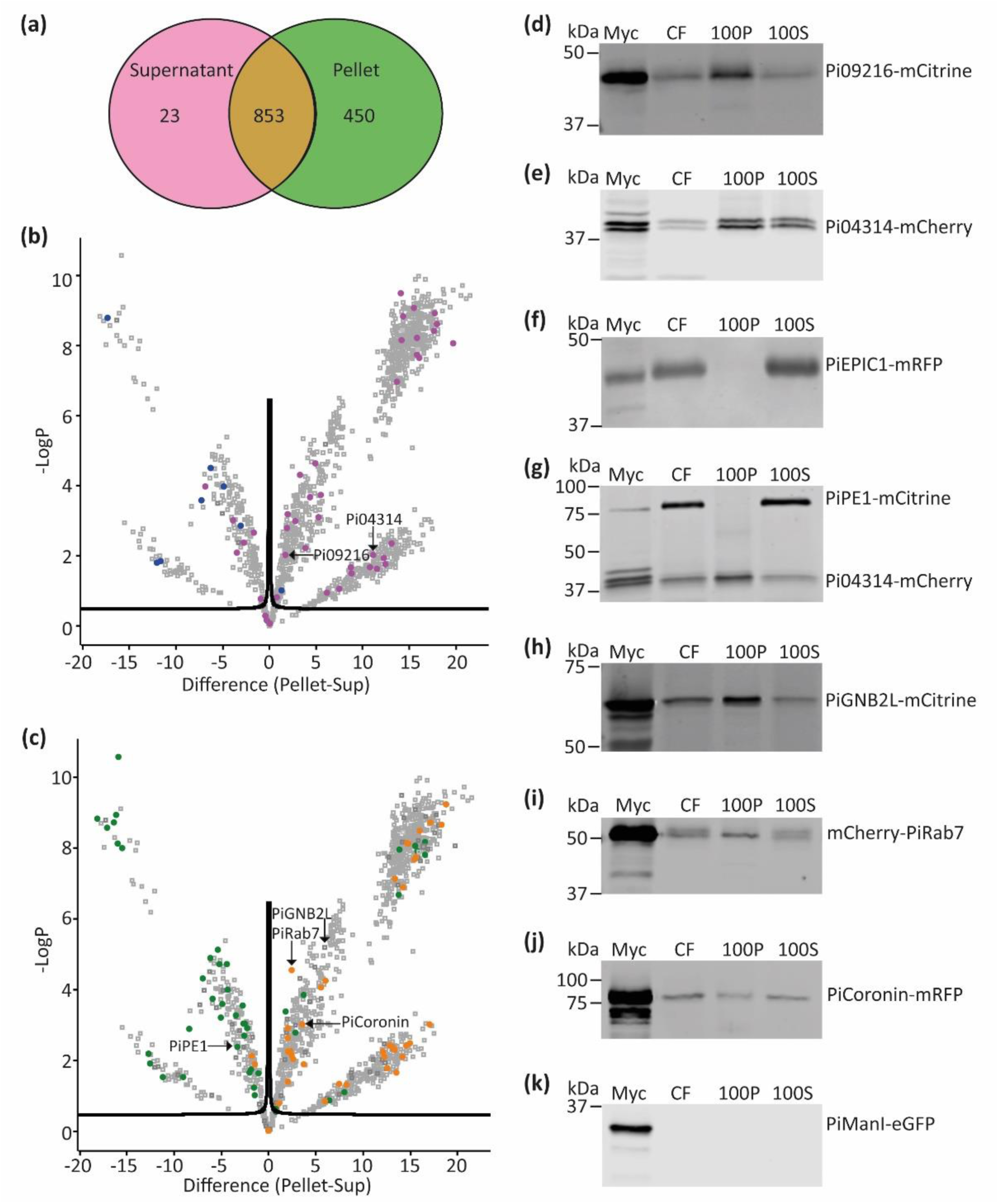
Overview of the *P. infestans* crude vesicle pellet proteome. (a) Venn diagram showing the distribution of proteins between the supernatant and pellet samples after crude extracellular vesicle pellet isolation and LC-MS/MS analysis from *P. infestans*-inoculated media. (b) Volcano plot showing the statistical distribution of proteins within the pellet vs supernatant samples. Pink spots indicate RxLR motif-containing effector proteins, blue spots represent apoplastic effector proteins. Black arrows highlight RxLR proteins of interest. (c) The same volcano plot with green spots indicating cell wall degrading enzymes (CWDE), orange spots indicating vesicle-associated proteins. Black arrows highlight CWDE and vesicle-associated proteins of interest. CWDE were identified using dbCAN3 (https://bcb.unl.edu/dbCAN2/). (d-k) Immunoblots of crude vesicle pellet and supernatant isolation from multiple transgenic *P. infestans* lines expressing mRFP, mCherry or mCitrine tagged proteins of interest. The membranes were probed with αRFP for mRFP or mCherry fusions, or αGFP for mCitrine fusions. These western blots show mycelia (Myc) and culture filtrate (CF) samples prior to any centrifugation, 100,000 x*g* pellet (100P) and supernatant samples after 100,000 x*g* spin (100S). These western blots confirm the distribution of the proteins based on the observations from the volcano plots (b-c). The proteins of interest expressed in the P. infestans isolates are (d) RxLR effector Pi09216-mCitrine. (e) RxLR effector Pi04314-mCherry. (f) apoplastic effector PiEPIC1-mRFP. (g) CWDE Pectinesterase 1 (PiPE1-mCitrine) and RxLR effector Pi04314-mCherry. (h) vesicle associated protein Guanine nucleotide-binding protein subunit β-2-like (PiGNB2L)-mCitrine. (i) vesicle associated protein PiRab7-mCherry. (j) vesicle associated protein PiCoronin-mRFP. (k) the control Golgi marker protein PiManI-eGFP (Ah-Fong and Judelson, 2011).

Perseus (version 2.0.6.0) was used to generate volcano plots determining which proteins were detected at significantly different amounts between the EV-associated pellet and EV-independent supernatant samples. Supplementary Figure S3a shows the volcano plot highlighting the different categories of proteins which were detected differentially. Group S1 indicates proteins that are found exclusively in the supernatant. S2 indicates proteins that are found in all supernatant samples and between 1 and 3 pellet samples but are statistically enriched within the supernatant. S3 indicates proteins that are found in all samples but are statistically enriched within the supernatant. Group P1 indicates proteins that are found exclusively in the pellet. P2 indicates proteins that are found in all pellet samples and between 1 and 3 supernatant samples but are statistically enriched within the pellet. P3 indicates proteins that are found in all samples but are statistically enriched within the pellet. Overall, 1009 proteins (76%) were enriched in the pellet, 255 proteins (19%) were enriched in the supernatant and 62 proteins (5%) were not significantly over-represented in either P or S samples (Table 1). 34% of proteins enriched in the supernatant were found to contain a signal peptide (SP), whereas only 8% of proteins enriched in the pellet had an SP (Table 1, Supplementary Table S3a).

**Table 1:** Enrichment of proteins of interest in the pellet vs the supernatant.

| Type of Protein | Non Significant | Pellet Enriched | Supt Enriched | Total |
| --- | --- | --- | --- | --- |
| All proteins | 62 | 1009 | 255 | 1326 |
| RxLRs | 3 | 33 | 6 | 42 |
| Apoplastic | 0 | 1 | 7 | 8 |
| Vesicle-associated | 1 | 38 | 2 | 41 |
| CWDE | 1 | 11 | 34 | 46 |
| TM | 2 | 71 | 8 | 81 |
| PAMPS | 1 | 4 | 8 | 13 |
| Signal Peptide | 11 | 63 | 81 | 155 |
| SP +TM | 0 | 22 | 6 | 28 |

Interestingly, the pellet was found to be significantly enriched in RxLR effector proteins (33 out of 42 proteins), suggesting that these effectors are potentially often associated with EVs (Figure 2b, Table 1, Supplementary Table S3b). By contrast, apoplastic effectors (7 out of 8 proteins) were predominantly detected in the supernatant samples showing that conventionally and non-conventionally secreted proteins may have different fates. It is worth noting that a number of RxLR (19) and apoplastic (9) effectors present in the original data (Supplementary Table S2a) were excluded from analyses as, being small proteins, they were only represented by a single peptide and were thus not included in the high confidence dataset (Supplementary Table S2b). Similar to the pattern for apoplastic effectors, secreted cell wall degrading enzymes (CWDEs) were predominantly detected in the supernatant (34 out of 46 proteins), whereas vesicle-associated proteins were predominantly enriched in the pellet samples (38 out of 41 proteins) (Figure 2c, Table 1, Supplementary Table S3b). PAMPs from *P. infestans* were found to be predominantly enriched in the supernatant samples (8 out of 13 proteins), although a few were enriched in the pellet (4 out of 13 proteins) (Supplementary Figure S3b, Table 1, Supplementary Table S3b).

To provide support for the observations in the proteomics dataset, transgenic *P. infestans* lines expressing a subset of genes of interest, tagged with fluorescent proteins, were subjected to the EV UC protocol and western blotting was used to investigate their presence in supernatant (S) and pellet (P) samples. We observed the enrichment of RxLR effectors Pi09216-mCitrine, Pi04314-mCherry and PiAVRblb1-mRFP in the pellet samples, whereas apoplastic effectors PiEPIC1-mRFP and PiSCR74-mRFP and the CWDE PiPE1-mCitrine were enriched in supernatant samples (Figure 2d-g, Supplementary Figure S3c,d). PiAVRblb1 and PiEPIC1, although only represented by single peptides were included, along with apoplastic effector SCR74 which was not detected in the proteome, as we had existing, well-characterised *P. infestans* transformants secreting these fusion proteins (Wang et al. 2017, 2018, 2023b). The pellet-to-supernatant ratio for three RxLRs vs three apoplastic effectors was quantified for multiple replicate western blots using LiCOR software and further supported patterns of enrichment in the proteomics data (Supplementary Figure S3e). A transformant co-expressing Pi04314-mCherry and PiPE1-mCitrine verified that these proteins were differentially enriched in pellet or supernatant samples, respectively (Figure 2g). *P. infestans* transformants expressing vesicle-associated proteins PiGNB2L-mCitrine and mCherry-PiRab7 also supported the proteomics data, whereas, although detected in the pellet, PiCoronin-mRFP was not enriched in these samples (Figure 2h-j). The control Golgi marker protein PiManl (Ah-Fong and Judelson 2011) was not detected in pellet or supernatant samples showing that not all proteins associated with cellular membrane compartments can be secreted or associated with EV-containing samples (Figure 2k).

### 3.3 MARVEL domain proteins are candidate EV markers

To identify potential EV marker proteins we sought transmembrane (TM)-containing proteins enriched in the pellet proteome. Unexpectedly, we did not identify any proteins corresponding to tetraspanins which are classic membrane markers of mammalian and plant EVs (Cai et al. 2021). However, we did identify 81 TM domain containing proteins and 71 of these (88 %) were significantly enriched in the pellet samples (Figure 3a, Table 1, Supplementary Table S3b). Of particular interest were two MARVEL domain-containing proteins, hereafter referred to as *P. infestans* MARVEL domain protein 1 and 2 (PiMDP1 and PiMDP2). A MARVEL domain consists of a bundle of four TM helices and PiPMDP1 and PiMDP2 each have two MARVEL domains, so contain a total of eight TM helices (Figure 3b) with the N- and C-termini predicted to be on the same side of the membrane. Interestingly yeast protein non classical export (NCE102) is a MARVEL domain protein which localises with the fungal EV marker SUR7 to eisosomes and is enriched in fungal EVs from *Candida albicans* (Dawson et al. 2020). This prompted us to examine PiMDP1 and PiMDP2 in more detail. Phylogenetic analysis using reciprocal best blast hit (RBBH) matches (Supplementary Figure S4a) showed that PiMDP1 and PiMDP2 candidate orthologues exist in all *Phytophthora* clades with PiMDP2 also present in many *Pythium* species indicating it is likely ancestral to PiMDP1. However, orthologues of these genes were not identified in the downy mildews as homology searches only yielded single MARVEL proteins that were not RBBH matches. Generally, each *Phytophthora* species has one copy of each *MDP* gene and these are arranged head-to-head with a small intervening region of several hundred base pairs (Supplementary Figure S4b), likely indicating a conserved evolutionary origin.

**Figure 3:**
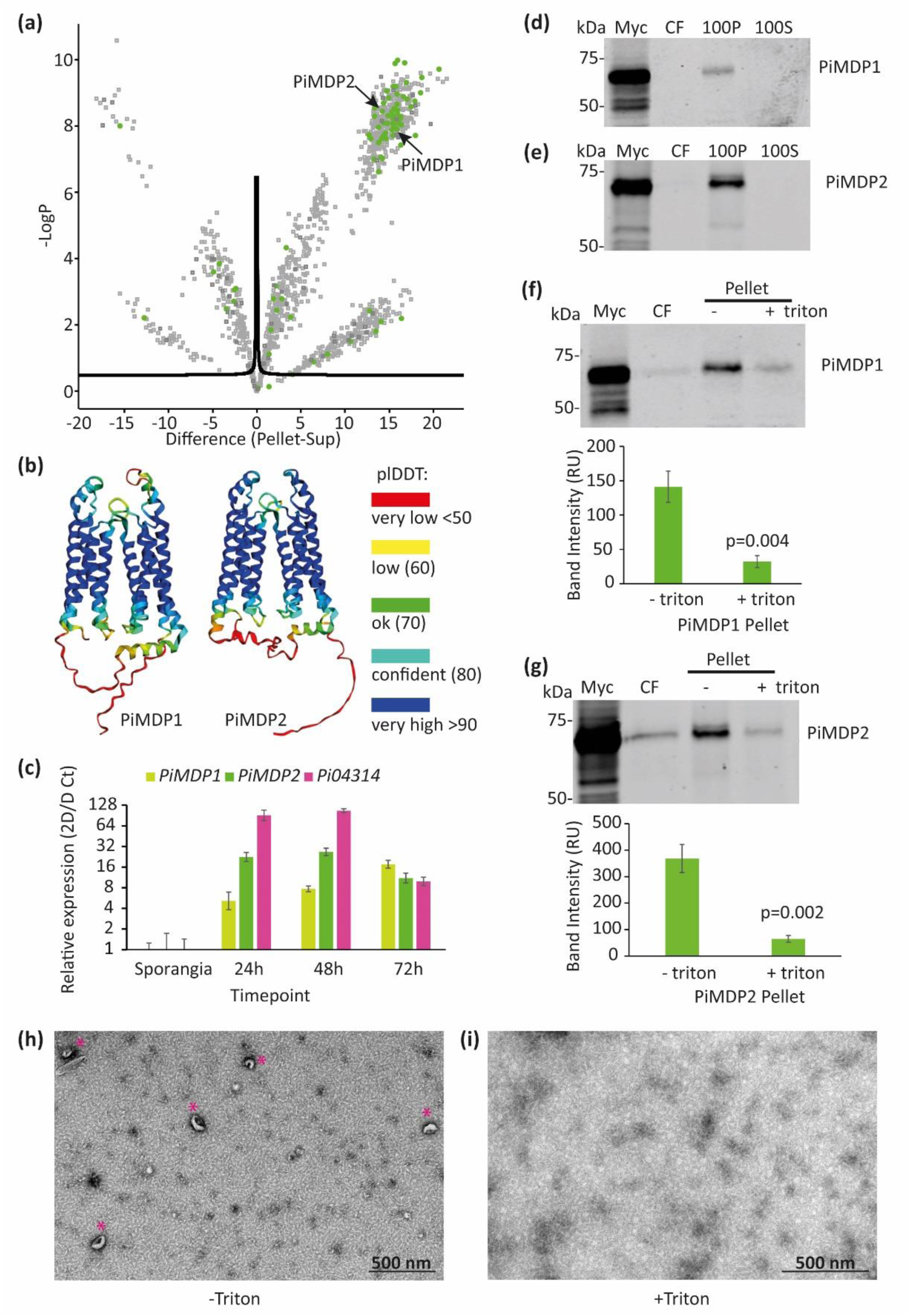
MARVEL-domain containing proteins are found exclusively in the crude vesicle pellet sample and are expressed during *P. infestans* infection. (a) Volcano plot showing the statistical distribution of proteins within the pellet vs supernatant samples. Green spots represent transmembrane (TM) domain containing proteins. Black arrows indicate the location of PiMDP1 and PiMDP2. TM domains were identified using DeepTMHMM (https://dtu.biolib.com/DeepTMHMM). (b) Predicted structures of PiMDP1 and PiMDP2 produced using Alphafold2. Protein structures are colour-coded to show plDDT confidence levels as shown in the key. (c) Gene expression of PiMDP1, PiMDP2 and the RxLR Pi04314 over a time-course of *P. infestans* isolate 3928A infecting Maris Piper potato leaves. Expression was normalised to the geometric mean of 3 housekeeping genes (ACTIN, Caesin kinase, Kelch domain repeat) using the 2D/D Ct method. Error bars show St Error. (d-e) crude vesicle pellet and supernatant isolation from transgenic *P. infestans* isolates expressing (d) PiMDP1-mCitrine tagged protein, (e) PiMDP2-mCitrine tagged protein. These western blots show mycelia (M) and culture filtrate (CF) samples prior to any centrifugation, 100,000 x*g* pellet (100P) and supernatant samples after 100,000 x*g* spin (100S). These western blots confirm the localisation of the proteins based on the observations from the volcano plot in (a). (f) -/+ 1% Triton treatment of culture filtrate from PiMDP1-mCitrine expressing *P. infestans* isolate. This western blot shows mycelia (M) and culture filtrate (CF) samples prior to any centrifugation and 100,000 x *g* pellet (100P) -/+ 1% Triton X-100. The band intensity graph shows the relative intensity of 100P -/+ 1% Triton bands from 6 independent replicates. Statistical analysis was done using a Paired T-test on SigmaPlot giving the two-tailed P value of 0.004. (g) -/+ 1% Triton treatment of culture filtrate from PiMDP2-mCitrine expressing *P. infestans* isolate. The western blot shows mycelia (M) and culture filtrate (CF) samples prior to any centrifugation and 100,000 x*g* pellet (100P) -/+ 1% Triton. The band intensity graph shows the relative intensity of 100P -/+ 1% Triton bands from 5 independent replicates. Statistical analysis was done using a Paired T-test on SigmaPlot giving the two-tailed P value of 0.002. (h) Transmission electron microscopy negative stain image showing vesicles present within a – Triton sample from a transgenic *P. infestans* isolate expressing both PiMDP1-mCitrine and Pi04314-mCherry. Asterisks indicate vesicles, scale bar represents 500 nm. (i) Transmission electron microscopy negative stain image indicating the loss of vesicles in an equivalent + Triton sample. Scale bar represents 500 nm.

The expression of *PiMDP1* and *PiMDP2* was examined during infection of Potato leaves with *P. infestans* and indicated that, while both genes are upregulated during infection (Figure 3c, Supplementary Figure S5a-b) *PiMDP1* transcript levels steadily increase throughout the infection time course, with the highest level at 72 hours post-inoculation (hpi). By contrast, *PiMDP2* expression is higher at the earlier timepoints of 24 and 48 hpi and thus more closely resembles the expression profile of RxLR effectors, such as *Pi04314,* which act as markers of the biotrophic phase of infection where *P. infestans* is coexisting with living plant tissue (Whisson et al. 2007).

To assess the suitability of PiMDP1 and PiMDP2 as candidate EV markers dual transgenic *P. infestans* lines were made co-expressing either PiMDP1-mCitrine or PiMDP2-mCitrine with RxLR effector Pi04314-mCherry. Initial EV ultracentrifugation preparations indicated that both PiMDP1 (Figure 3d) and PiMDP2 (Figure 3e) accumulate in the pellet and not the supernatant of the 100,000 g spin. To provide further evidence that these proteins are associated with EVs the culture filtrate was equally divided and one half was treated with detergent 1% Triton X-100 prior to the 100,000 x g spin. Figure 3f shows that in the presence of Triton significantly less PiMDP1 is recovered as detected by western blotting and quantified using the LICOR quantification system. The same result was observed with PiMDP2 (Figure 3g). TEM analysis of pellets without Triton showed the presence of EV-like structures (Figure 3h) whereas in the presence of Triton no EVs were observed (Figure 3i). These results suggest that these candidate markers are associated with membranous EV-like structures which are sensitive to disruption with detergents such as Triton. As PiMDP1 and 2 are predicted to be integral membrane proteins IP experiments were carried out to determine if the termini of these proteins are present within or exposed on the surface of the vesicular structures. Supplementary Figure S6 shows that C-terminally tagged PiMDP1-mCitrine and PiMDP2-mCitrine are only immunocaptured in the presence of the detergent triton suggesting that the tagged ends are inside the EVs, and thus not exposed until vesicles are disrupted (Supplementary Figure S6a-b). The same result is observed when each of the N-termini are tagged with mCherry (Supplementary Figure S6c-d).

To further characterise these marker proteins the crude pellets of the transgenic *P. infestans* lines were subjected to further separation and purification on sucrose density columns. Firstly, a transformant co-expressing two RxLRs, Pi09216-mCitrine and Pi04314-mCherry (Figure 4a-b, Supplementary Figure S7a-b), showed each effector fusion to accumulate predominantly in Fraction 4 with a sucrose density of ∼1.16 to 1.18 g/cm^2^. MARVEL marker candidate PiMDP1-mCitrine and RxLR Pi04314-mCherry were observed also to predominantly co-accumulate in sucrose density Fraction 4 (Figure 4c-d, Supplementary Figure S7c-d) as did PiMDP2-mCitrine and Pi04314-mCherry co-expressed in transgenic *P. infestans* (Figure 4e-f, Supplementary Figure S7e-f). Moreover, a transgenic line co-expressing both MARVEL domain proteins demonstrated that mCherry-PiMDP1 and PiMDP2-mCitrine both accumulated primarily in Fraction 4 (Figure 4g-h, Supplementary Figure S7g-h). Fraction 4 correlates with the density that EVs such as exosomes are expected to be buoyant (1.13-1.19 g/ml), according to the literature (Colombo et al. 2014). Nanoparticle tracking analysis (NTA) of the sucrose fractions indicated that the largest concentration of nanoparticles appeared in Fraction 4 with a main peak sizes of approximately 80 nm diameter and a smaller peak of approximately 180 nm (Figure 4i). Few nanoparticles were observed in Fractions 1 and 2; Fraction 3 had a small peak around 140nm and there was a low level mixture of larger sized material in Fractions 5 and 6. In agreement with this, TEM images showed no EVs in Fraction 1 (Figure 4j), as a control, whereas many EV-like vesicle structures were observed in Fraction 4 samples (Figure 4k).

**Figure 4:**
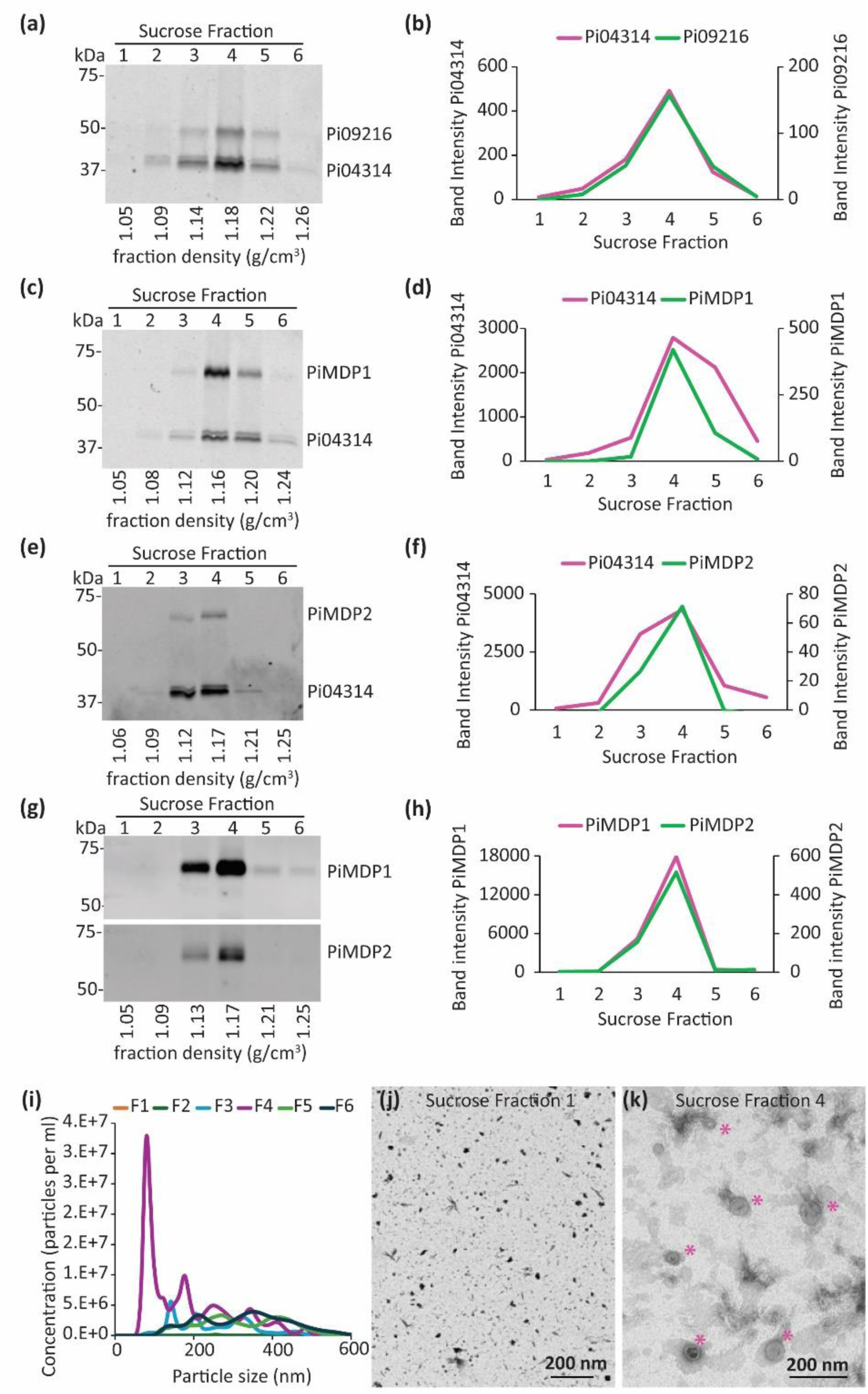
MARVEL-domain and RxLR effector proteins are detected within the same sucrose density fraction. Crude vesicle pellets were top-loaded onto a discontinuous sucrose gradient consisting of 10, 20, 30, 40, 50, 60 and 70% layers. After centrifugation at 100K x*g* for 16 h, six fractions of 1.9 ml each were collected and processed with further ultracentrifugation to obtain a pure vesicle pellet. All membranes were dual probed with αGFP and αRFP. The density (g/cm^3^) of each fraction is detailed under each blot. (a) Six fractions isolated using a top-loaded crude pellet from a P. infestans isolate expressing both Pi09216-mCitrine and Pi04314-mCherry. (b) Graph showing band intensity for Pi09216-mCitrine (green) and Pi04314-mCherry (pink) from each fraction shown in (a) (c) Fractions isolated from *P. infestans* expressing both PiMDP1-mCitrine and Pi04314-mCherry. (d) Graph showing band intensity for PiMDP1-mCitrine (green) and Pi04314-mCherry (pink) from each fraction shown in (c). (e) Fractions isolated from *P. infestans* expressing both PiMDP2-mCitrine and Pi04314-mCherry. (f) Graph showing band intensity for PiMDP2-mCitrine (green) and Pi04314-mCherry (pink) from each fraction shown in (e). (g) Fractions isolated from *P. infestans* expressing both PiMDP2-mCitrine and mCherry-PiMDP1. (h) Graph showing band intensity for PiMDP2-mCitrine (green) and mCherry-PiMDP1 (pink) from each fraction shown in (g). (i) Nanoparticle tracking (NTA) data showing the number and size distribution of particles in each sucrose fraction. Results show the average from three independent replicates for each sample. (j) Transmission electron microscopy negative stain image of a sample of fraction 1 isolated from *P. infestans* isolate expressing both PiMDP1-mCitrine and Pi04314-mCherry. (k) Transmission electron microscopy negative stain image showing a range of vesicles in fraction 4 isolated from the same isolate. Asterisks indicate EV’s, scale bars in I and j represent 200 nm.

### 3.4 MARVEL domain proteins co-localise frequently with RxLR Pi04314 at hyphal vesicles *in vitro*

After establishing that the MARVEL EV marker candidates co-accumulate with RxLR Pi04314 in Sucrose Fraction 4 that has been demonstrated to contain EV structures by NTA and TEM, the localisation of these proteins inside *P. infestans* hyphae was examined to determine whether they also co-associated with vesicles prior to secretion. Firstly, hyphae from a transgenic line co-expressing Pi04314-mCherry and the apoplastic effector/CWDE PiPE1-mCitrine (Wang et al 2018), which does not accumulate in the EV pellet (Figure 2g), were studied by confocal microscopy. As expected for two secreted proteins, both localised to vesicle-like structures inside the hyphae but fluorescence intensity plots drawn through the vesicles showed little co-localisation of the two proteins (Figure 5a, 5f, Supplementary Figure S8a, S9). A transgenic line co-expressing two independent RxLRs, Pi04314-mCherry and Pi09216-mCitrine, was examined and both effectors strongly co-localised to the same vesicle-like structures inside the hyphae, suggesting that they may follow the same secretory pathway (Figure 5b, 5f, Supplementary Figure S8b, S9). EV marker candidates PiMDP1-mCitrine and PiMDP2-mCitrine were both observed to localise to the plasma membrane (PM) as well as to many vesicle-like structures inside the hyphae (Figure 5c, 5d, Supplementary Figure S8c-e). Fluorescence intensity plots through the hyphae demonstrate PiMDP1-mCitrine and Pi04314-mCherry showed the same strong levels of colocalisation to vesicles as the two RXLR effectors (Figure 5, Supplementary Figure S8c, S9). PiMDP2-mCitrine and Pi04314-mCherry also co-localised to a subset of vesicles, albeit to lower levels than PiMDP1-mCitrine and Pi04314-mCherry (Figure 5, Supplementary Figure S8d, S9). In addition, when mCherry-PiMDP1 and PiMDP2-mCitrine were co-expressed *in vitro* both localise to subsets of overlapping and distinct vesicle-like structures in hyphae (Figure 5, Supplementary Figure S8e, S9). Together, this suggests that there is a mixed population of vesicles containing diverse cargoes that may have different functions and distinctive transit routes. To further confirm the identity of the vesicle-like structures observed in the hyphae endo-vesicles were isolated from *in vitro* grown hyphae expressing mCherry-PiMDP1 and PiMDP2-mCitrine (Reynolds et al. 2014; Wang et al. 2023b). The resulting pelleted endo-vesicles were found to contain the tagged proteins by immunoblotting (Supplementary Figure S10a-b) and were observed to be sensitive to the detergent triton, suggesting that the PiMDP1 and PiMDP2 markers are indeed incorporated in membranous vesicles within hyphae.

**Figure 5:**
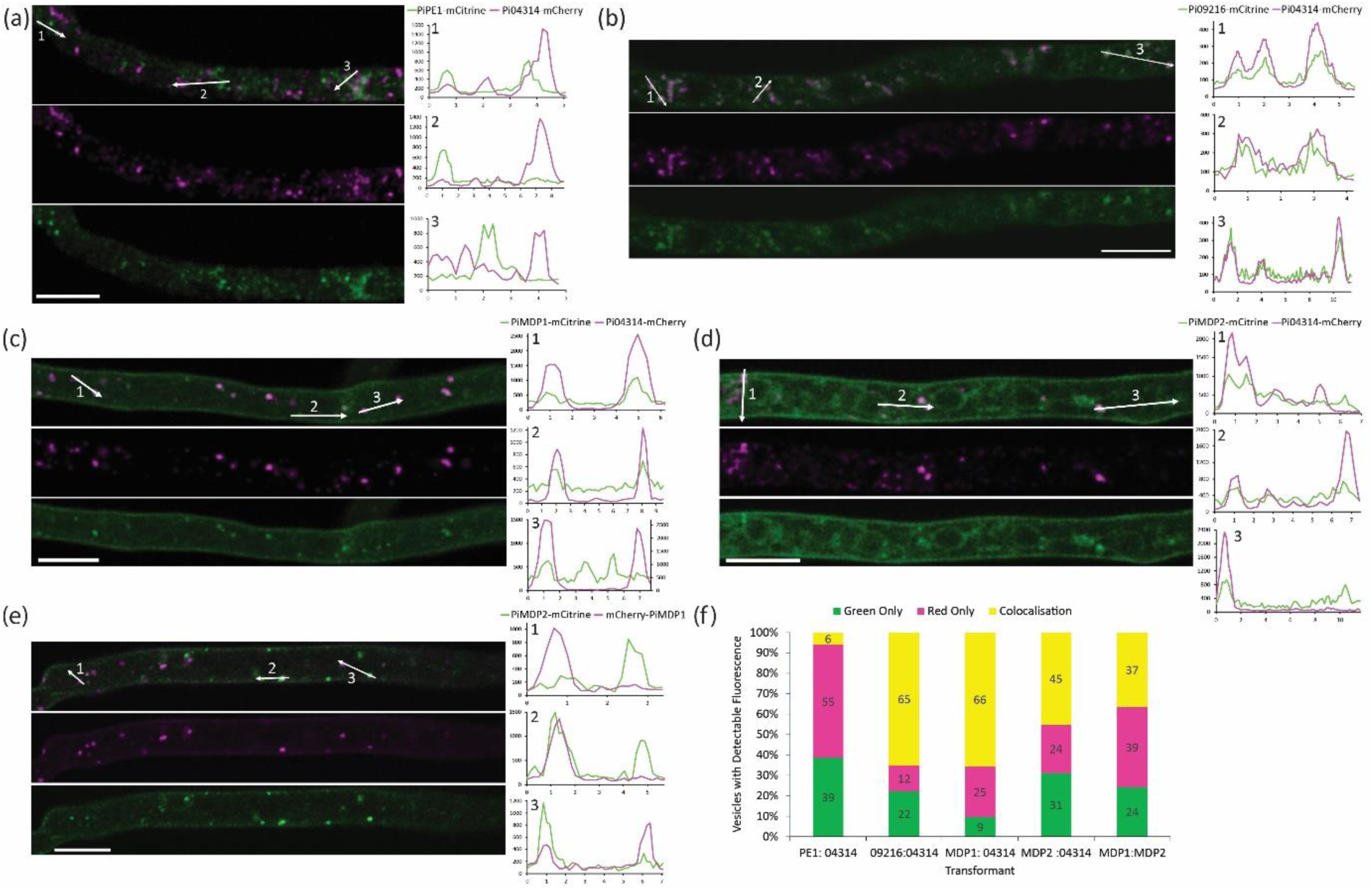
PiMDP1/PiMDP2 and RxLR proteins colocalise in vesicle-like structures in hyphae grown *in vitro*. Projection images from confocal z series images of hyphae from transformed *P. infestans* transgenic lines expressing fluorescent protein fusions grown on microscope slides. The mCitrine (green) and mCherry (magenta) channels are shown separately and merged with three numbered arrows indicating transects drawn using single optical sections to produce the three numbered fluorescence intensity plots to the right. The transgenic lines were co-expressing: (a) apoplastic effector PiPE1-mCitrine and RxLR Pi04314-mCherry; (b) RxLRs Pi09216-mCitrine and Pi04314-mCherry; (c) PiMDP1-mCitrine which localises to the PM and also partially co-localises to vesicles with Pi04314-mCherry; (d) PiMDP2-mCitrine which localises to the PM and also partially co-localises to vesicles with Pi04314-mCherry; (e) PiMDP2-mCitrine which partially co-localises to vesicles with mCherry-PiMDP1; both are observed at the PM. Scale bars are 10 µm. Fluorescence intensity plots show relative fluorescence on the y axis against distance in µm on the x axis. (f) Stacked bar chart shows quantification of the percentage of vesicles showing mCitrine only, mCherry only or colocalisation of both fluorophores. Results are the average counts from at least 12 images; percentages are shown on the chart; n indicates the total number of vesicles counted for each transformant (PiPE1-mCitrine & Pi04314-mCherry n= 612; Pi09216-mCitrine & Pi04314-mCherry n= 350; PiMDP1-mCitrine & Pi04314-mCherry n= 237; PiMDP2-mCitrine & Pi04314-mCherry n= 303; PiMDP2-mCitrine & mCherry-PiMDP1 n= 293).

### 3.5 MARVEL-domain and RxLR effector proteins co-localise frequently at vesicles during early infection

As both MARVEL domain EV marker candidates were found to partially co-localise with RxLR effector Pi04314 in vesicles inside *in vitro* grown hyphae, we examined their localisation during early plant infection. To investigate this, we imaged hyphae just behind the leading edge of infection, where haustoria were just starting to form and are represented by small protrusions on the hyphae where RxLRs accumulate. Haustoria are specialised infection structures produced by the pathogen and are the sites in closest association with the host. These structures act as major sites of secretion for both pathogen and host (Wang et al. 2017, 2018; Boevink et al. 2020; Bozkurt and Kamoun 2020). In all cases Pi04314-mCherry was observed to accumulate around the base of developing haustoria (Figure 6a-b, Supplementary Figure S11) as has been reported in previous publications (Wang et al. 2017, 2018). Both PiMDP1-mCitrine and PiMDP2-mCitrine were localised at the PM, including in the developing haustorial membrane (HM) (Figure 6a-b, Supplementary Figure S11). In addition, fluorescence intensity plots through the hyphae show that PiMDP1-mCitrine and Pi04314-mCherry (Figure 6a, Supplementary Figure S11a) and PiMDP2-mCitrine and Pi04314-mCherry (Figure 6b, Supplementary Figure S11b) were observed to colocalise to a subset of vesicle-like structures inside the hyphae but not every vesicle was found to contain both proteins, as was shown for *P. infestans* hyphae *in vitro*. This agrees with the observations within *in vitro* grown hyphae, providing further evidence of both shared and distinct vesicle trafficking and potential downstream functions.

**Figure 6:**
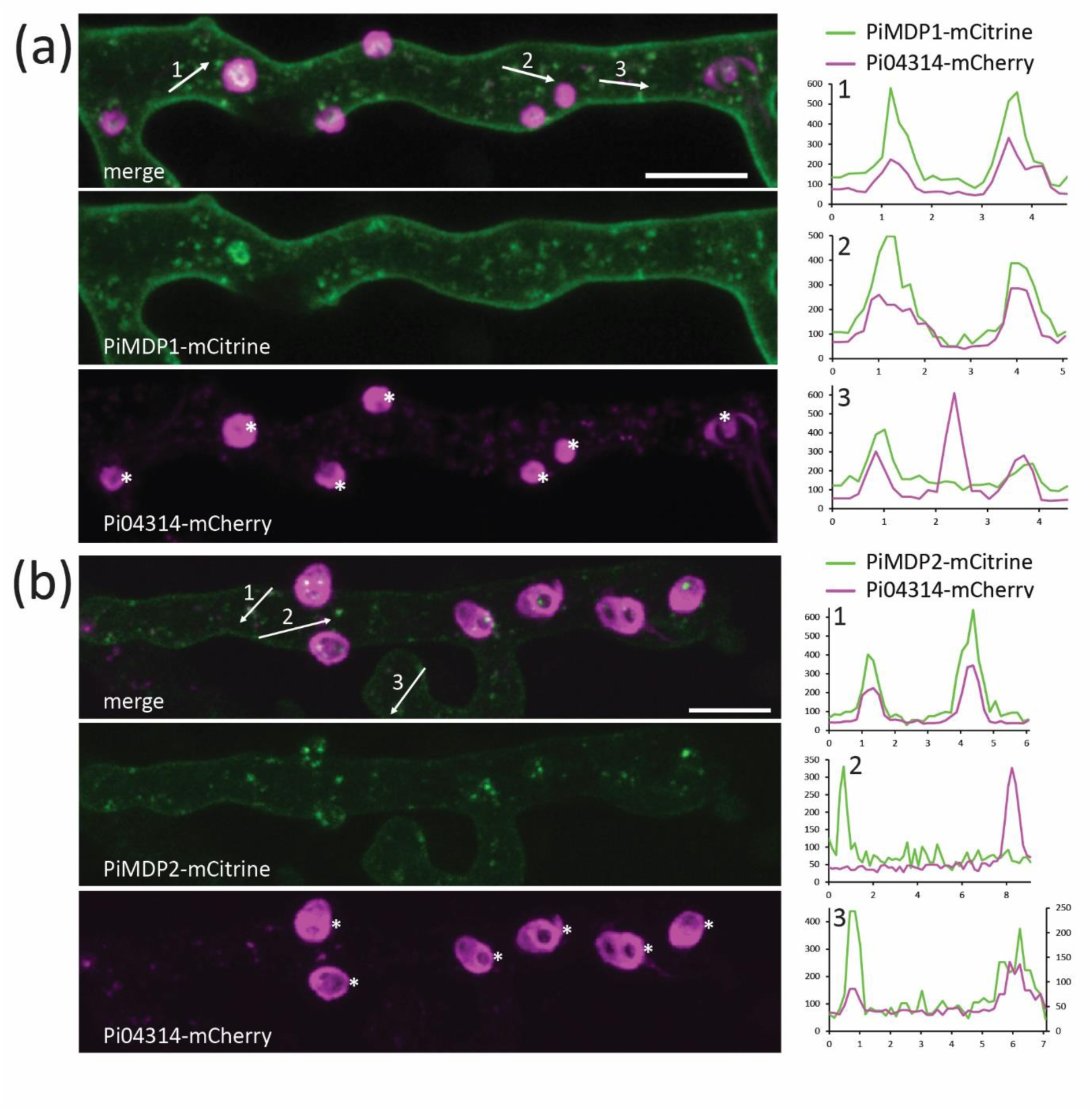
PiMDP and RxLR proteins co-localise in vesicle-like structures in hyphae during infection. Projection images from confocal z series of hyphae from transformed *P. infestans* isolates expressing fluorescent protein fusions infecting *N. benthamiana* leaves. The mCitrine (green) and mCherry (pink) channels are shown separately and merged with three numbered arrows indicating transects drawn using single optical sections to produce the three numbered fluorescence intensity plots below. (a) PiMDP1-mCitrine localises to the plasma membrane (PM) and partially co-localises to vesicles with Pi04314-mCherry. (b) PiMDP2-mCitrine localises to the PM and partially co-localises to vesicles with Pi04314-mCherry. The mCherry intensities were low in this hypha and thus are plotted with the right-hand vertical axis to make the patterns more obvious. Asterisks indicate haustoria and scale bars are 10 µm. Fluorescence intensity plots show relative fluorescence on the y axis against distance in µm on the x axis.

### 3.6 PiMDP2 is enriched in the mature haustorial membrane during infection

Later stages of infection, where the haustoria on *P. infestans* hyphae were mature and formed elongated hook-like projections, were then examine by confocal microscopy. Here a striking difference in the MARVEL-domain candidate EV marker localisation was observed. PiMDP1-mCitrine was localised as before to the PM, HM and vesicles inside the hyphae (Figure 7a, Supplementary Figure S12a). However, PiMDP2-mCitrine was found to accumulate strongly at the HM (Figure 7b, Supplementary Figure S12b), although it was still observed weakly in the PM and in vesicles. In both cases the co-expressed Pi04314-mCherry was found to be localised around the base of the haustoria as expected. The difference in PiMDP2 localisation was quantified by measuring the fluorescence intensity ratio of both PiMDP1-mCitrine and PiMDP2-mCitrine in the HM relative to the PM in multiple images. While PiMDP1-mCitrine found equally present in both membranes with a ratio of approximately 1, the fluorescence of PiMDP2-mCitrine was significantly stronger in the HM with a higher mean ratio (Figure 7c). This points to some possible differences in function between our EV candidate markers.

**Figure 7:**
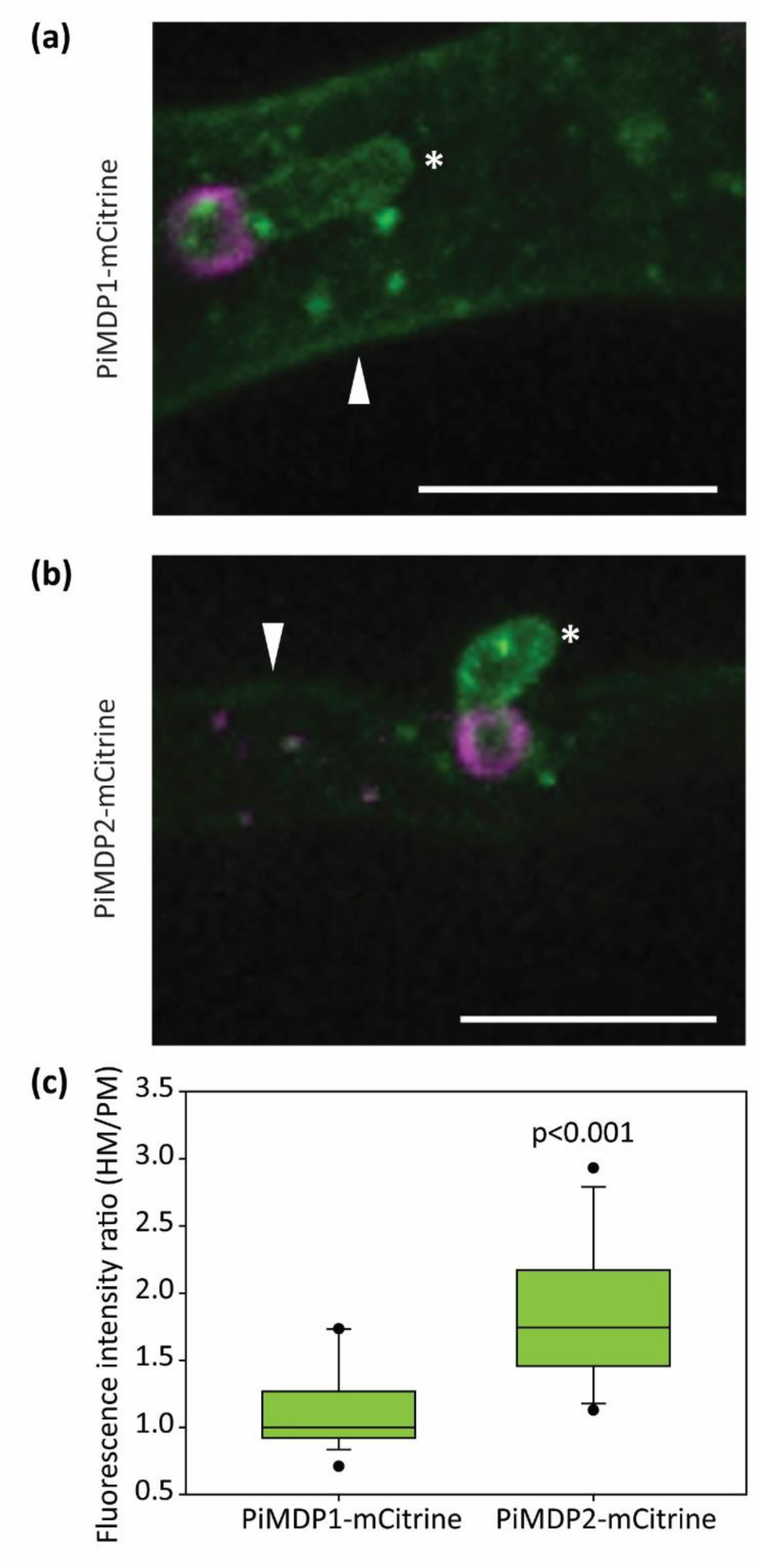
PiMDP2-mCitrine accumulates in the mature haustorial membrane during infection. Confocal projection images from z series images of *P. infestans* hyphae and haustoria during plant infection showing (a) PiMDP1-mCitrine localised to the plasma membrane (PM), haustorial membrane (HM) and vesicle-like bodies in the hypha, with 04314-mCherry localised around the haustoria and, (b) PiMDP2-mCitrine accumulates more strongly in the HM with little PM fluorescence, in addition to vesicle-like bodies in the hypha, with 04314-mCherry localised around the haustoria. Asterisks indicate haustoria, arrowheads indicate the PM and the scale bars are 10 μM. (c) Boxplot shows quantification of the ratio of the relative fluorescence intensity in regions of the HM compared to the PM measured from single optical sections. The points indicate the 5^th^ and 95^th^ percentile outliers and the lines show the median values. Statistical analysis was done using a Mann-Whitney Rank Sum Test on SigmaPlot giving the P value of <0.001, (n≥ 19).

## 4.0 DISCUSSION

Differential ultracentrifugation is a common technique used to isolate EVs from different organisms and tissues. Here, the pellet from ultracentrifugation (UC) of the culture media used to grow *P. infestans* mycelia *in vitro* was demonstrated to contain EVs by both NTA and TEM analyses (Figure 1). Proteomic analysis of the pelleted (EV-associated) material following UC showed that it is enriched for both vesicle-associated and transmembrane proteins compared to the supernatant (EV-independent) samples (Table 1), which is consistent with the pellet containing EVs. Moreover, the pellets also contain proteins from filamentous plant pathogens, such as coronin, which acts to regulate actin organisation and vesicle trafficking during infection (Dulal et al. 2021; Li et al. 2019), as well as Rab7, a GTPase which regulates effector secretion and membrane trafficking (Chen et al. 2023). Proteins which are notable in their absence in the *P. infestans* EV-associated proteome are tetraspanins, which are typically enriched in EVs in mammalian and plant systems (Cai et al. 2021; Van Niel et al. 2018). Tetraspanins contain four TM helices and are used as molecular markers of EVs. They are abundant, present in many different cell types and form stable tetraspanin-associated microdomains. The availability of many monoclonal tetraspanin antibodies allows their purification by immunoprecipitation thus facilitating studies of EV biogenesis, trafficking and analysis of EV cargoes (Wang et al. 2024). The presence of EV markers is crucial to bring EV research in oomycetes to the cutting edge.

Interestingly, tetraspanin homologues were also not identified in proteomic studies of EVs in some fungi, so other molecular markers have been proposed. One such marker is SUR7 which was found to be enriched in EVs from *Candida albicans* and *Zymoseptoria tritici* (Dawson et al. 2020; Hill and Solomon 2020). SUR7 is a TM protein containing four TM helices and is a component of the Membrane Compartment of Can1 (MCC)/eisosome, a specialised microdomain at the fungal PM which forms stable membrane furrows (Lanze et al. 2020). *SUR7* knockouts in *C. albicans* were shown to form smaller EVs with altered cargoes containing less fungal virulence proteins (McKenna et al. 2023). The MCC/eisosome is involved in many different membrane processes, including membrane integrity, lipid regulation, maintaining nutrient transporters at the PM, and responses to stress, as well as impacting fungal virulence (Lanze et al. 2020). Another component of the MCC/eisosome is Non Classical Export (NCE)102 which was also found to be enriched in fungal EVs (Dawson et al. 2020). In addition to being part of the MCC/eisosome, NCE102 is reportedly involved in unconventional protein secretion in yeast (Cleves et al. 1996). NCE102 is a <u>M</u>AL <u>A</u>nd <u>R</u>elated proteins for <u>VE</u>sicle formation and membrane <u>L</u>ink (MARVEL) domain-containing protein. A MARVEL domain consists of four TM helices and the best characterised mammalian family members are involved in membrane trafficking, transcytosis and cell signalling (Sánchez-Pulido et al. 2002; Li et al. 2024). Plant pathogenic fungus *Alternaria brassicicola* in which AbNCE102 was mutated had a reduced ability to colonise its host *B. oleracea,* possibly due to the significant reduction in the formation of appressoria-like structures required for invasion (Colou et al. 2019). Like NCE102, PiMDP1 and PiMDP2 are PM-localised and are enriched in EVs. However, they possess two MARVEL domains and so contain eight TM helices. Human MARVEL protein MYADM also contains two MARVEL domains, is PM localised alongside the Rho GTPase Rac and is involved in membrane organisation, cell spreading and migration (Aranda et al. 2011; Labat-de-Hoz et al. 2023). Moreover, MYADM acts as a receptor required for entry of human parechovirus PeV-A (Qiao et al. 2024) and has been proposed as a molecular marker of EVs from plasma (Hu et al. 2024).

Although tetraspanins were not observed in our EV proteome, they were observed in a reported EV proteome for *Phytophthora sojae* (Zhu et al 2023). It is not clear why tetraspanins were absent in our EV dataset, although growth conditions were different. However, of the 50 most abundant non-ribosomal proteins in our EV proteome (Supplementary Table S2), candidate orthologues of 19 (38%) were represented in the *P. sojae* EV proteome (Supplementary Table S4). Importantly, the *P. sojae* orthologue of PiMDP1 (Ps_341763) was represented in the *P. sojae* EV proteome (Zhu et al 2023), indicating that it is a potential EV marker also for that pathogen.

### The Role of Unconventional Protein Secretion (UPS) in Filamentous Plant Pathogens

The secretion of EVs is known to be one form of UPS which normally occurs for proteins which do not contain a signal peptide (SP). This study used proteomic analysis to identify proteins secreted into the culture media used to grow *P. infestans*. Of the proteins identified as enriched in the supernatant 35% were predicted to contain an SP which included many apoplastic effectors and CWDE known to be conventionally secreted. Only 8% of proteins found to be enriched in the pellet contained an SP, many of which included RxLR effectors which have been reported to be unconventionally secreted (Wang et al. 2018, 2017; Xu 2023). This suggests that UPS contributes significantly to the *P. infestans* secretome. Indeed the EV pellet proteome exhibits some of the hallmarks of UPS and includes proteins such as ATG9 which is associated with autophagosomes but also with Compartments for UPS (CUPS) (Bruns et al. 2011; New and Thomas 2019). The pellet also contains the vacuolar sorting receptor BP-80, which is associated with multivesicular bodies (MVBs) that have been implicated in the biogenesis of EVs which play a role in plant pathogen interactions (Zhuang et al. 2024; Cai et al. 2021). UPS has been suggested to play an important role in filamentous plant pathogens with a large proportion of secreted proteins found to lack an SP in *Magnaporthe*, *Fusarium*, *Trichoderma* and *Phytophthora*. Moreover, increases of UPS in these pathogens have been reported during abiotic stress and notably during infection (Stuer et al. 2023).

### Multiple Secretory Pathways

RxLR Pi04314 and apoplastic effector/CWDE pectinesterase 1 (PiPE1) were observed to localise to distinct vesicles inside plant hyphae suggesting that they are exported by different secretion pathways, in line with previous observations (Wang et al. 2018). However, co-expression of the RxLR effectors Pi04314 and Pi09216 revealed them to display predominant co-localisation to the same vesicles suggesting a common secretory pathway (Figure 5, Supplementary Figure S8a-b, S9). In the case of candidate EV markers PiMDP1 and PiMDP2 there is frequent but not complete co-localisation at vesicles in hyphae *in vitro* and *in planta* with the RXLR Pi04314 and, indeed, with each other (Figures 5-6, Supplementary Figures S8 & S9, S11). This makes a case for the existence of both distinct and overlapping pathways for vesicle export. It is also possible that Pi04314 is associated with different vesicles prior to its association with PiMDP-marked vesicles. The fact that both PiMDP1 and PiMDP2 are predominantly found, alongside Pi04314, in sucrose density fractions indicative of EV accumulation (1.13-1.19 g/ml; Colombo et al 2014), where we observe many vesicular structures by TEM and NTA, plus both markers are vulnerable to treatment with detergent (Figures 3, 4, Supplementary Figure S10), suggests that they are indeed associated with membranous structures such as EVs. Interestingly during *in planta* colonisation there are signs of diverging functional roles for PiMDP1 and PiMDP2, as the latter shares a gene expression pattern with virulence factors such as Pi04314 (Figure 3c; Supplementary Figure S5) and is thus associated with the biotrophic phase of *P. infestans* infection, when both pathogen and host are interacting closely via the haustorial interface. By contrast, *PiMDP1* gene expression steadily increases throughout infection from biotrophy into necrotrophy. Moreover, PiMDP2 but not PiMDP1 accumulates strongly in the mature haustorial membrane (HM) (Figure 7, Supplementary Figure S12). Such enrichment of PiMDP2 at the HM could be consistent with its accumulation as a result of vesicle fusion during secretion.

A previous study has shown that RxLRs can be taken up by plants via clathrin-mediated endocytosis (CME) (Wang et al. 2023b, 2023a). Indeed, there is some overlap (34%) in the RxLRs identified in the proteomics of immune-purified endosomes and those identified in this study in the EV pellet proteome (Supplementary Table S5). One question is how is such endocytosis triggered? The recognition and binding of PAMPs by their cognate receptors, such as the binding of flg22 by FLS2, is known to induce CME of the receptor (Mbengue et al. 2016). The majority of PAMPs identified in this study were abundantly represented in the supernatant. However, some were enriched in the EV pellet. Further work is needed to determine whether such PAMPs are exposed on the surface of EVs and so trigger uptake through binding to pattern recognition receptors on the host PM. Interestingly, although our study did not reveal tetraspanins (TETs), TETs associated with *P. sojae* EVs were found to be recognised as PAMPs by host and nonhost plants (Zhu et al. 2023).

### EVs Associated with Virulence Factors

It is attractive to hypothesise that secretion of proteins such as virulence factors may be best achieved by coordinating their delivery to precise locations through their packaging into EVs. 42 RXLR effector proteins were detected in our *in vitro* study of the EV proteome. This is remarkable, given that the genes encoding these virulence proteins are transcriptionally up-regulated during infection (e.g. Whisson et al 2007). However, one explanation for their detection is that, despite the likely low basal expression level of many of these RXLR effectors, enriching for an EV pellet co-enriched 33 RXLRs to detectable levels. Future work will hopefully enable pathogen EVs to be immuno-isolated from infection, using tagged marker proteins such as PiMDPs revealed in this study. This will allow us to better determine the protein cargoes and composition of EVs during infection.

Consistent with our study of *in vitro* EVs from *P. infestans*, proteomic analysis of EVs from the plant pathogen *Fusarium graminearum* identified a collection of both existing and novel candidate effector proteins and virulence factors associated with EVs (Garcia-Ceron et al. 2021). Infiltration of plants with EVs from *F. oxysporum* f. sp. *vasinfectum* led to host cell death (Bleackley et al. 2020). Also in oomycete research, *P. capsici* EVs enhanced virulence when applied to leaves (Fang et al. 2021). Identifying molecular markers associated with EVs, such as PiMDP1 and PiMDP2, especially those EVs also associated with virulence factors, will be important to dissect secretion pathways required for infection. Such markers also provide novel agrochemical control targets. This study provides a large dataset of EV pellet-associated proteins which can be dissected by the community to determine if useful markers for EV biogenesis, distinct secretion pathways, and thus targets for disease control, can be identified.

## Acknowledgements

We thank Prof Simon Powis from the University of St Andrews for his expertise and use of Nanoparticle tracking equipment. We thank also Prof Alex Jones, Warwick University, for helpful guidance in analysing proteomic data.

## Author contributions

S.B., H.M., W.W., S.C.W., P.C.B., and P.R.J.B. conceived of and designed the research; H.M., S.B., W.W., P.C.B., L.W., J.P., SW., and S.C.W performed the research; S.B., H.M., W.W., J.P., S.W., and S.C.W. contributed new reagents/analytic tools; S.B., H.M., P.C.B., S.C.W., W.W., and P.R.J.B. analysed data; P.R.J.B., S.C.W. and P.C.B. won funding for the research; and H.M., S.B, P.C.B and P.R.J.B. wrote the paper with input from all co-authors.

The authors declare no competing interest.

