## Supplementary Figure for "Identification of MARVELlous Protein Markers for *Phytophthora infestans* Extracellular Vesicles"

**Supplementary Figures**

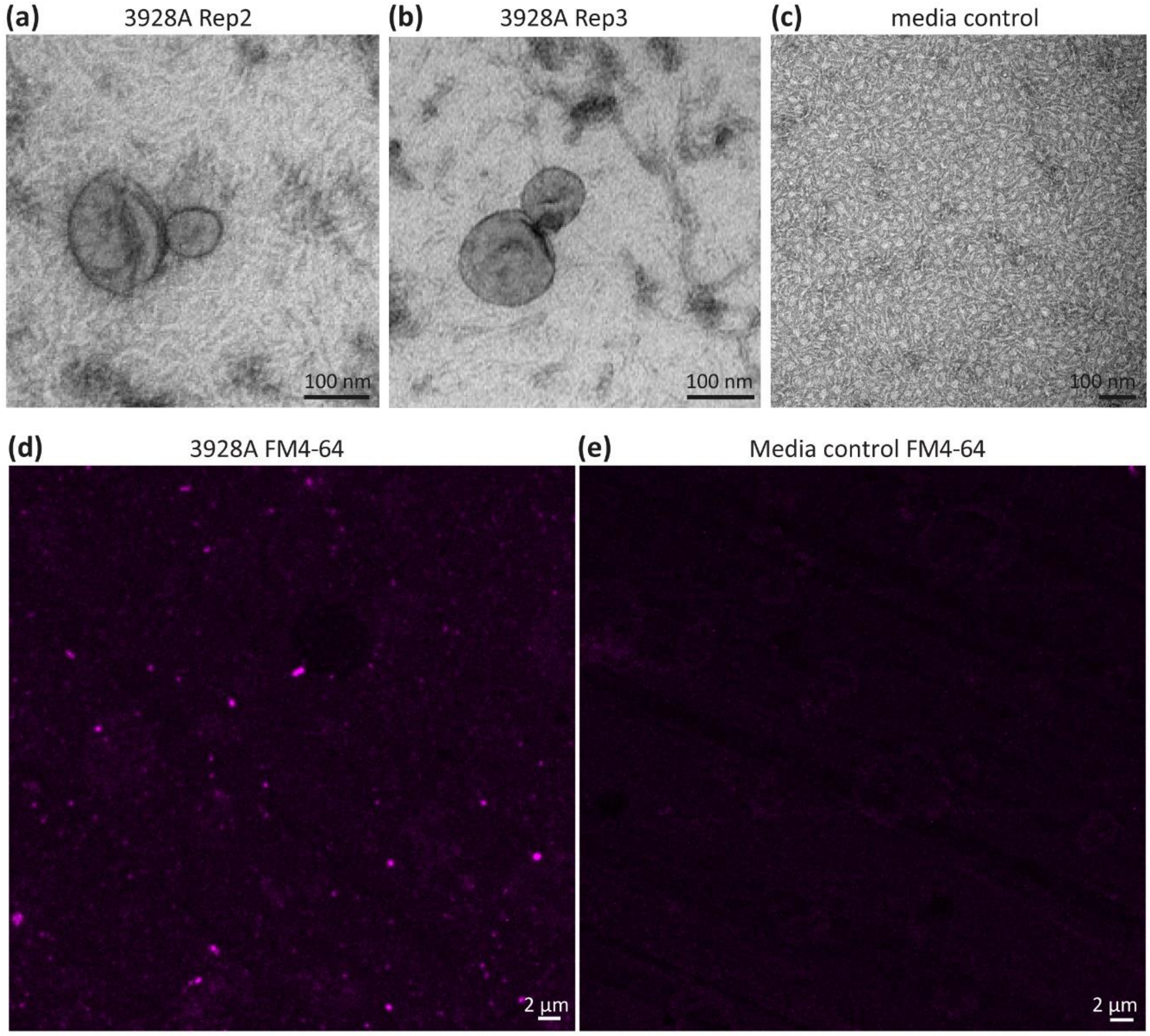


**Figure S1: Extracellular vesicles identified in crude pellets.** (a-b) Transmission electron microscopy negative stain images showing EVs present within crude nanoparticle/vesicle pellets isolated from media from *P. infestans* strain 3928A cultures. (c) Transmission electron microscopy negative stain image of the crude nanoparticle pellet isolated from uninoculated media. Scale bar represents 100 nm. (d) Confocal z series projection of FM4-64 staining of crude nanoparticle/vesicle pellets isolated from media from *P. infestans* strain 3928A. (e) Confocal z series projection of FM4-64 staining of crude pellets isolated from uninoculated media, scale bar is 2 µm

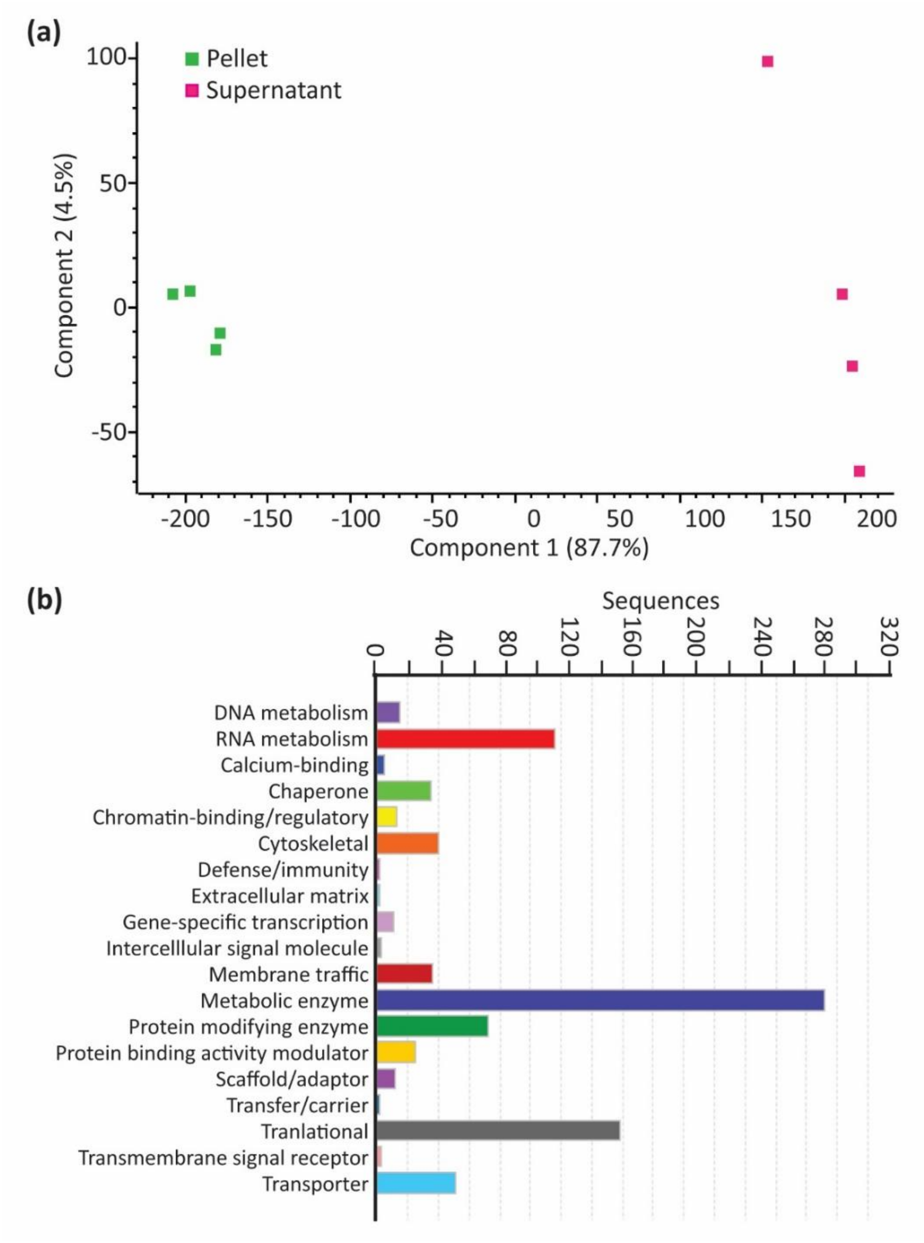


**Figure S2: Analysis of the *P. infestans* crude vesicle pellet proteome.** (a) Graph of principal component analysis (PCA) of proteomics results. Pink squares represent the four replicates of *P. infestans* crude vesicle pellet samples while the green squares represent the four replicates of supernatant from the *P. infestans* inoculated samples. (b) Graph of GO term enrichment of proteins identified from the *P. infestans* crude vesicle pellet samples.

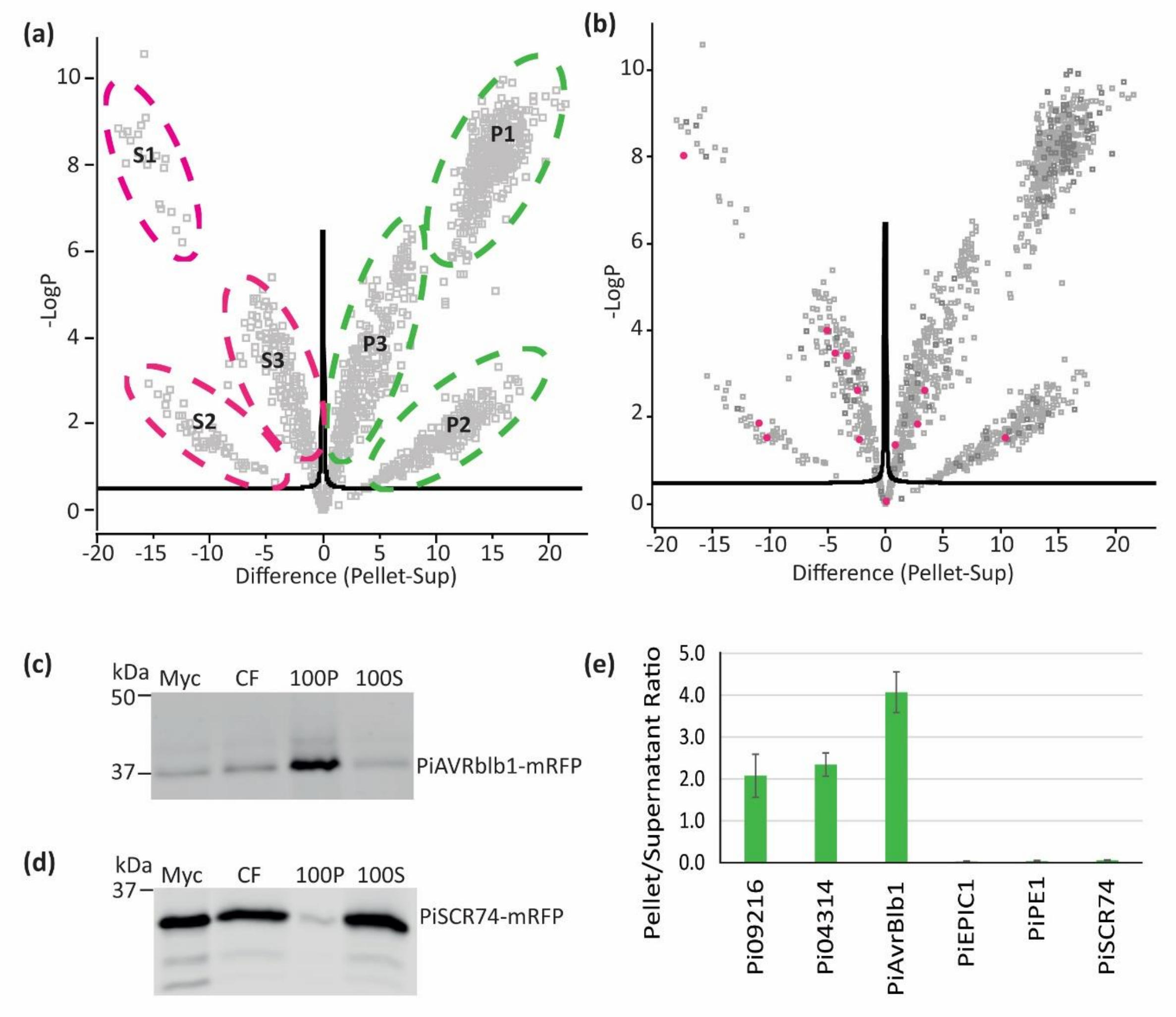


**Figure S3: Overview of the *P. infestans* crude vesicle pellet proteome.** (a) Volcano plot showing the 3 categories of proteins identified. S1 indicates proteins that are found exclusively in the supernatant in all four replicates. S2 indicates proteins that are found in all four supernatant samples and up to 3 of the pellet samples but are statistically enriched within the supernatant. S3 indicates proteins that are found in all four supernatant and pellet samples but are statistically enriched within the supernatant. P1 indicates proteins that are found exclusively in the pellet in all four replicates. P2 indicates proteins that are found in all four pellet samples and up to 3 of the supernatant samples but are statistically enriched within the pellet. P3 indicates proteins that are found in all four pellet and supernatant samples but are statistically enriched within the pellet. (b) Volcano plot showing the statistical distribution of proteins within the pellet vs supernatant samples. Pink spots represent PAMPs. (c-d) Crude vesicle pellet and supernatant isolation from two transgenic *P. infestans* lines expressing mRFP tagged proteins. These western blots show mycelia (Myc) and culture filtrate (CF) samples prior to any centrifugation, 100,000 × g pellet (100P) and supernatant samples after 100,000 × g spin (100S). (c) *P. infestans* line expressing RxLR effector PiAVRblb1-mRFP. (d) *P. infestans* line expressing apoplastic effector PiSCR74-mRFP. (e) A graph showing the band intensity ratio between pellet vs supernatant protein bands for Pi09216-mCitrine, Pi04314-mCherry, PiAVRblb1-mRFP, PiEPIC1-mRFP, PiPE1-mCitrine and PiSCR74-mRFP. Band intensity data is from a minimum of 3 biological replicates.

(a)

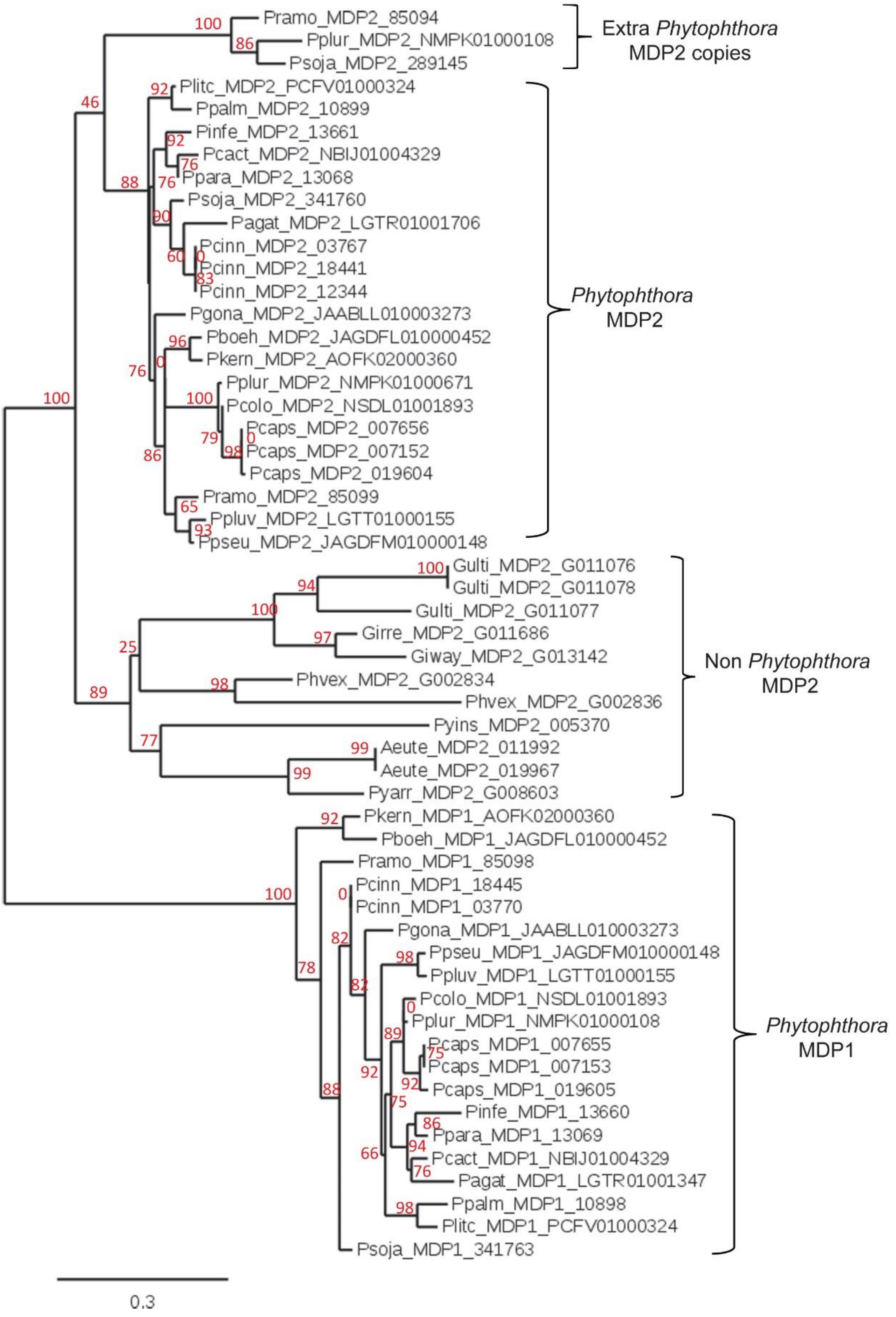
Pramo_MDP2_85094
Pplur_MDP2_NMPK01000108
Psoja_MDP2_289145
Extra *Phytophthora* MDP2 copies
Plitc_MDP2_PCFV01000324
Ppalm_MDP2_10899
Pinfe_MDP2_13661
Pcact_MDP2_NBIJ01004329
Ppara_MDP2_13068
Psoja_MDP2_341760
Pagat_MDP2_LGTR01001706
Pcinn_MDP2_03767
Pcinn_MDP2_18441
Pcinn_MDP2_12344
Pgona_MDP2_JAABLL010003273
Pboeh_MDP2_JAGDFL010000452
Pkern_MDP2_AOFK02000360
Pplur_MDP2_NMPK01000671
Pcolo_MDP2_NSDL01001893
Pcaps_MDP2_007656
Pcaps_MDP2_007152
Pcaps_MDP2_019604
Pramo_MDP2_85099
Ppluv_MDP2_LGTT01000155
Ppseu_MDP2_JAGDFM010000148
*Phytophthora* MDP2
Gulti_MDP2_G011076
Gulti_MDP2_G011078
Gulti_MDP2_G011077
Girre_MDP2_G011686
Giway_MDP2_G013142
Phvex_MDP2_G002834
Phvex_MDP2_G002836
Pyins_MDP2_005370
Aeute_MDP2_011992
Aeute_MDP2_019967
Pyarr_MDP2_G008603
Non *Phytophthora* MDP2
Pkern_MDP1_AOFK02000360
Pboeh_MDP1_JAGDFL010000452
Pramo_MDP1_85098
Pcinn_MDP1_18445
Pcinn_MDP1_03770
Pgona_MDP1_JAABLL010003273
Ppseu_MDP1_JAGDFM010000148
Ppluv_MDP1_LGTT01000155
Pcolo_MDP1_NSDL01001893
Pplur_MDP1_NMPK01000108
Pcaps_MDP1_007655
Pcaps_MDP1_007153
Pcaps_MDP1_019605
Pinfe_MDP1_13660
Ppara_MDP1_13069
Pcact_MDP1_NBIJ01004329
Pagat_MDP1_LGTR01001347
Ppalm_MDP1_10898
Plitc_MDP1_PCFV01000324
Psoja_MDP1_341763
*Phytophthora* MDP1
0.3

(b)

| Oomycete Species | Gene copy number | | Space between MDP genes | Gene Orientation |
|---|---|---|---|---|
| | MPD1 | MDP2 | | |
| *P. infestans* | 1 | 1 | 387 bp | → ← |
| *P. capsici* | 3 | 3 | 1287, 87, 236 bp | → ←, → ←, → ← |
| *P. cinnamomi* | 2 | 3 | 7986, 7960 bp (other genes in between) | → ←, → ← |
| *P. palmivora* | 2 | 2 | 222 bp | → ← |
| *P. parasitica* | 1 | 2 | 58 bp | → ← |
| *P. ramorum* | 1 | 2 | 366 bp | → ← |
| *P. sojae* | 1 | 2 | 4632 bp (other genes in between, incl. another MDP2) | → ← |
| *P. plurivora* | 1 | 2 | 265 bp | → ← |
| *P. colocasiae* | 1 | 1 | 249 bp | → ← |
| *P. cactorum* | 1 | 1 | 260 bp | → ← |
| *P. pseudosyringae* | 1 | 1 | 401 bp | → ← |
| *P. pluvialis* | 1 | 1 | 415 bp | → ← |
| *P. kernoviae* | 1 | 1 | 1701 bp | → ← |
| *P. boehmeriae* | 1 | 1 | 1621 bp | → ← |
| *P. gonapodyides* | 1 | 1 | 139 bp | → ← |
| *P. agathidicida* | 1 | 1 | | |
| *P. litchii* | 1 | 1 | 257 bp | → ← |
| *Phytopythium vexans* | 0 | 2 | | |
| *Globisporangium irregulare* | 0 | 1 | | |
| *Globisporangium iwayamae* | 0 | 1 | | |
| *Globisporangium ultimum* | 0 | 3 | | |
| *Pythium aphanidermatum* | 0 | 1 | | |
| *Pythium arrhenomanes* | 0 | 1 | | |
| *Pythium insidiosum* | 0 | 1 | | |
| *Aphanomyces euteiches* | 0 | 4 | | |

**Figure S4: Phylogenetic analysis of Marvel domain containing proteins within Oomycetes.** (a) Phylogenetic neighbour-joining tree of full length MDP proteins from oomycetes. Orthologs were inferred by reciprocal best blast match (RBBM). The multiple sequence alignment was undertaken in Clustal omega, and the tree was generated using Phylogeny.fr (http://www.phylogeny.fr/) (Dereeper et al., 2008). This program uses an approximation of the standard likelihood ratio test instead of bootstrapping.Values are shown in red. Pinfe = *P. infestans*, Pcaps = *P. capsici*, Pcinn = *P. cinnamomi*, Ppalm = *P. palmivora*, Ppara = *P. parasitica*, Psoja = *P. sojae*, Pramo = *P. ramorum*, Pplur = *P. plurivora*, Pcolo = *P. colocasiae*, Ppluv = *P. pluvialis*, Ppseu = *P. pseudosyringae*, Pkern = *P. kernoviae*, Pboeh = *P. boehmeriae*, Pagat = *P. agathidicida*, Plitc = *P. litchi*, Pcact = *P. cactorum*, Pgona = *P. gonapodyides*, Girre = *Globisporangium irregulare*, Giway = *G. iwayamae*, Gulti = *G. ultimum*, Phvex = *Phytopythium vexans*, Pyins = *Pythium insidiosum*, Pyarr = *Py. arrhenomanes*, Aeuti = *Aphanomyces euteiches*. (b) A table detailing information on MDP orthologues from oomycetes including species they have been identified within, copy number, distance between genes in the genome and gene orientation.

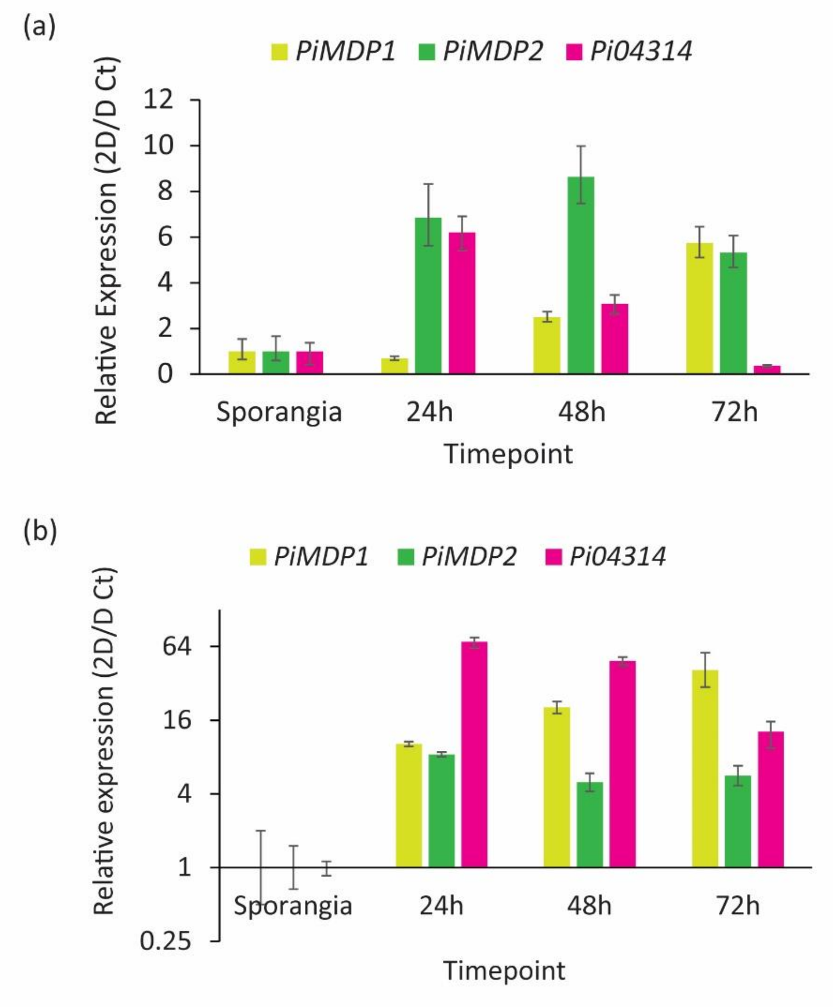


**Figure S5: MARVEL domain and *Pi04314* gene expression over infection time course.** Gene expression of *PiMDP1, PiMDP2* and the RxLR *Pi04314* over an infection time course of *P. infestans* isolate 3928A on potato cultivar Maris Piper leaves. Each graph represents an independent biological replicate. Expression was normalised to the geometric mean of 3 housekeeping genes (ACTIN, Caesin kinase, Kelch domain repeat) using the 2D/D Ct method. Error bars show Standard Error.

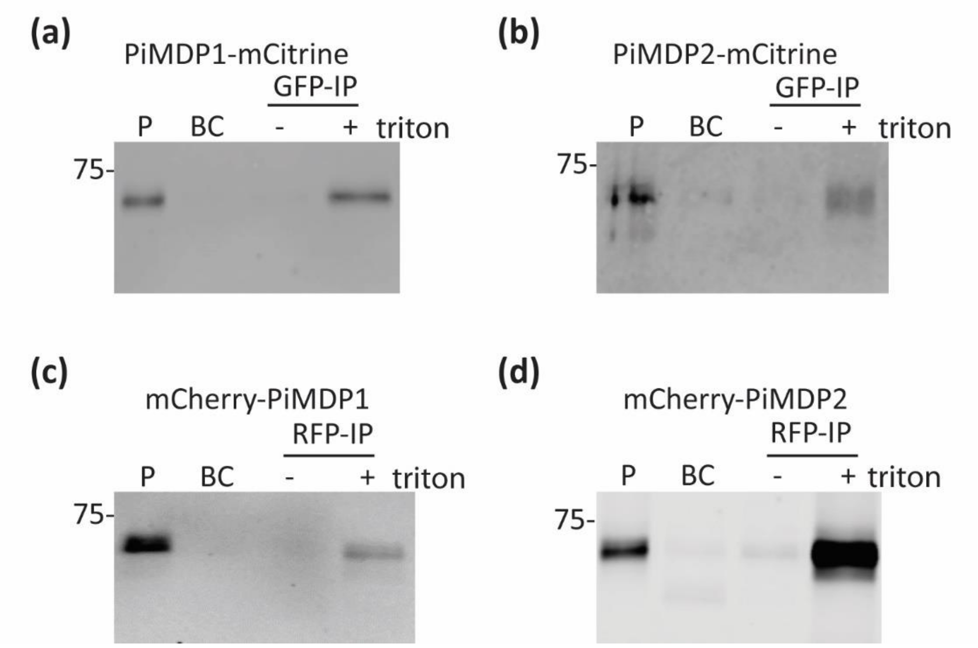


**Figure S6: PiMDP1 and PiMDP2 are only immunocaptured in the presence of Triton.** Crude pellets (P) were isolated by ultracentrifugation and resuspended in 1 × PBS, this sample was incubated with binding control (BC) beads to demonstrate no non-specific binding, samples were split and treated with (+) or without (-) Triton x100 detergent before incubation with the appropriated GFP or RFP trap magnetic beads. Blots show that PiMDP1 and PiMDP2 were only immunocaptured in the presence of detergent treatment showing that the tagged ends of the proteins are located inside the membrane structures. PiMDP1 (a) and PiMDP2 (b) were tagged independently with a C-terminal mCitrine tag and PiMDP1 (c) and PiMDP2 (d) were also tagged with N-terminal mCherry tags with the same results observed. Size markers are kDa.

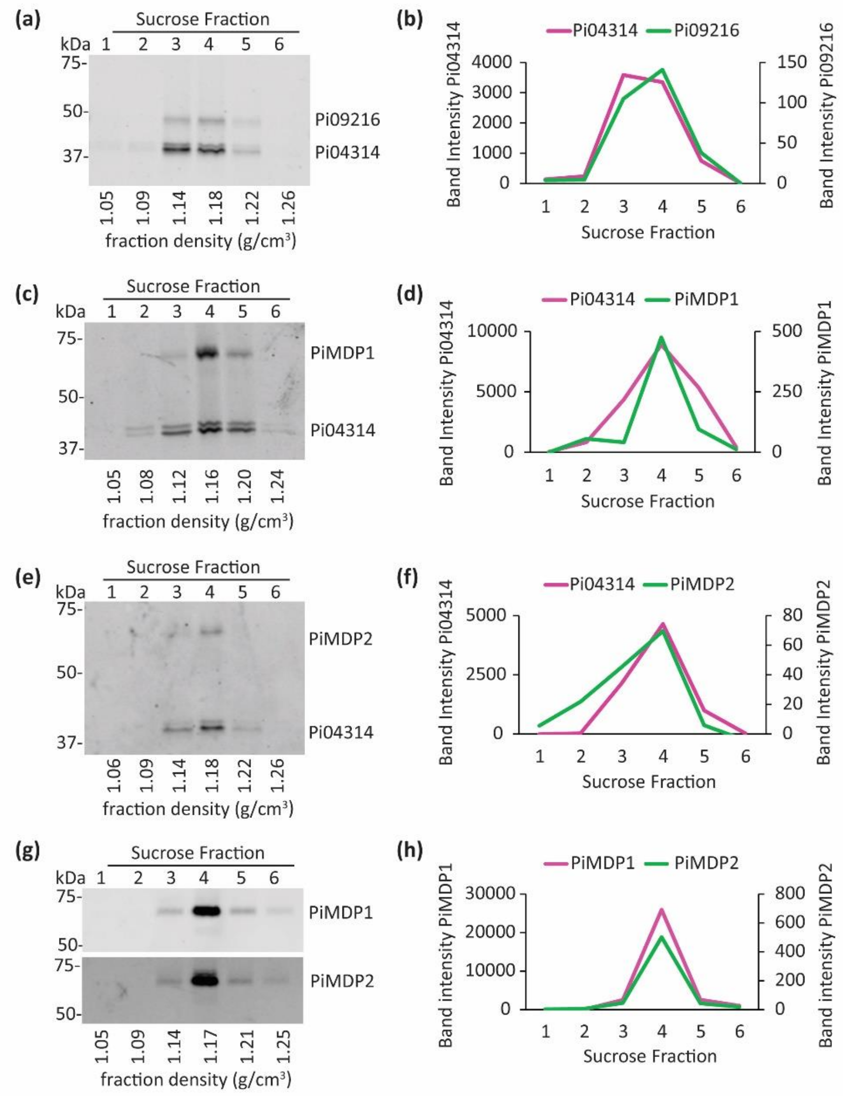


**Figure S7: MARVEL-domain and RxLR effector proteins detected in the same sucrose density fraction.** Independent biological replicates compared to Fig 4 showing consistency of experimentation. Crude vesicle pellets were top-loaded onto a discontinuous sucrose gradient consisting of 10, 20, 30, 40, 50, 60 and 70% layers. After centrifugation at 100,000 × g for 16 h, six fractions of 1.9 ml each were collected and processed with further ultracentrifugation to obtain a vesicle pellet representing each fraction. All membranes were dual probed with αGFP and αRFP. The density (g/cm$^3$) of each fraction is detailed under each blot. (a) Six fractions from a *P. infestans* isolate expressing both Pi09216-mCitrine and Pi04314-mCherry. (b) Graph showing band intensity for Pi09216-mCitrine (green) and Pi04314-mCherry (pink) from each fraction shown in (a). (c) Six fractions isolated using a top-loaded crude pellet from a *P. infestans* isolate expressing both PiMDP1-mCitrine and Pi04314-mCherry. (d) Graph showing band intensity for PiMDP1-mCitrine (green) and Pi04314-mCherry (pink) from each fraction shown in (c). (e) Six fractions isolated using a top-loaded crude pellet from a *P. infestans* isolate expressing both PiMDP2-mCitrine and Pi04314-mCherry. (f) Graph showing band intensity for PiMDP2-mCitrine (green) and Pi04314-mCherry (pink) from each fraction shown in (e). (g) Six fractions isolated using a top-loaded crude pellet from a *P. infestans* isolate expressing both PiPiMDP2-mCitrine and mCherry-PiMDP1. (h) Graph showing band intensity for PiMDP2-mCitrine (green) and mCherry-PiMDP1 (pink) from each fraction shown in (g).

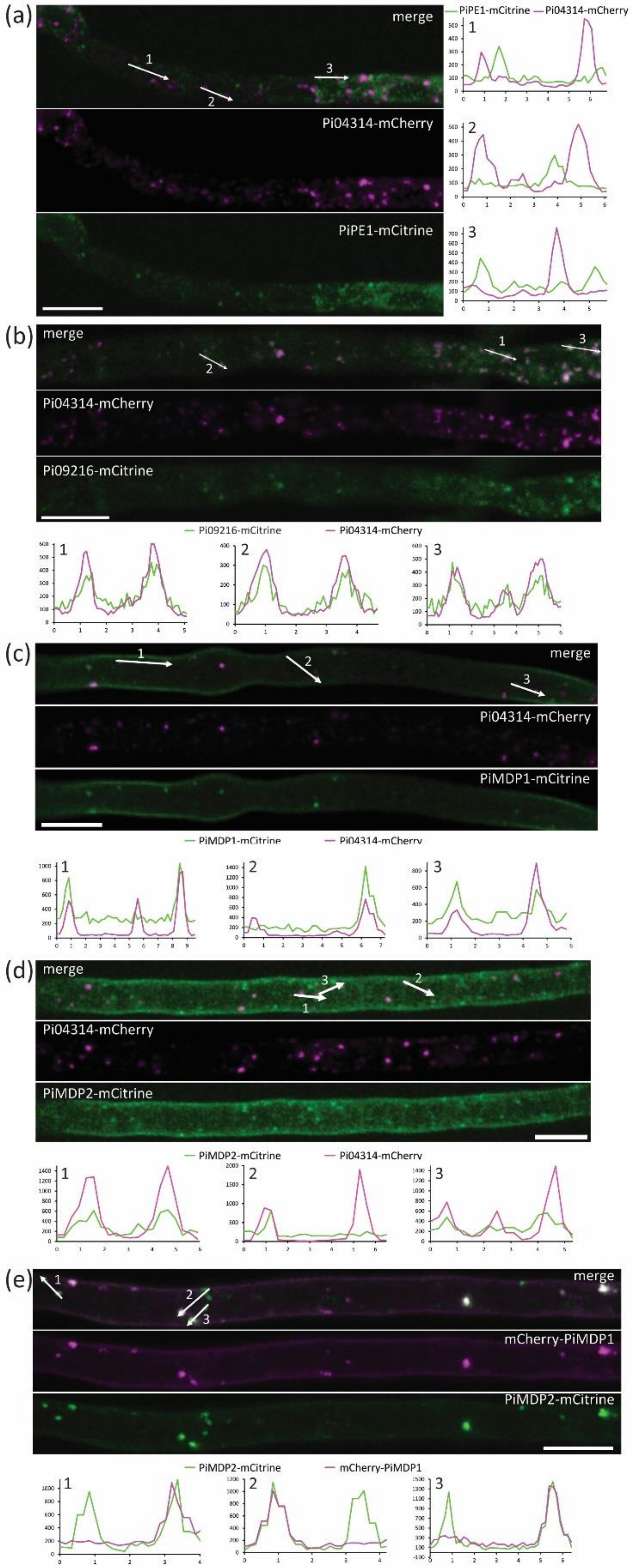

(a)
merge
Pi04314-mCherry
PiPE1-mCitrine
PiPE1-mCitrine
Pi04314-mCherry
(b)
merge
Pi04314-mCherry
Pi09216-mCitrine
Pi09216-mCitrine
Pi04314-mCherry
(c)
merge
Pi04314-mCherry
PiMDP1-mCitrine
PiMDP1-mCitrine
Pi04314-mCherry
(d)
merge
Pi04314-mCherry
PiMDP2-mCitrine
PiMDP2-mCitrine
Pi04314-mCherry
(e)
merge
mCherry-PiMDP1
PiMDP2-mCitrine
PiMDP2-mCitrine
mCherry-PiMDP1

**Figure S8: MARVEL-domain and RxLR effector proteins colocalise in vesicle-like structures in hyphae grown in vitro.** An additional example of results as shown in Figure 5. Images are projections of confocal z series of transformants grown on microscope slides showing the mCitrine (green) and mCherry (pink) channels and merged images with three numbered arrows indicating example transects drawn on single optical sections to produce the three numbered fluorescence intensity plots below. (a) Apoplastic effector PiPE1-mCitrine co-expressed with RxLR Pi04314-mCherry, plots indicate no colocalization. (b) Co-expression of two RxLRs; Pi09216-mCitrine with Pi04314-mCherry. Intensity plots show strong colocalization. (c) PiMDP1-mCitrine localises to the PM and partially co-localises at vesicle-like bodies with Pi04314-mCherry. (d) PiMDP2-mCitrine localises to the PM and partially co-localises at vesicles with Pi04314-mCherry. (e) PiMDP2-mCitrine partially co-localises at vesicles with mCherry-PiMDP1, both are observed in the PM. Scale bars are 10 µm. Fluorescence intensity plots show relative fluorescence on the y axis against distance in µm on the x axis.

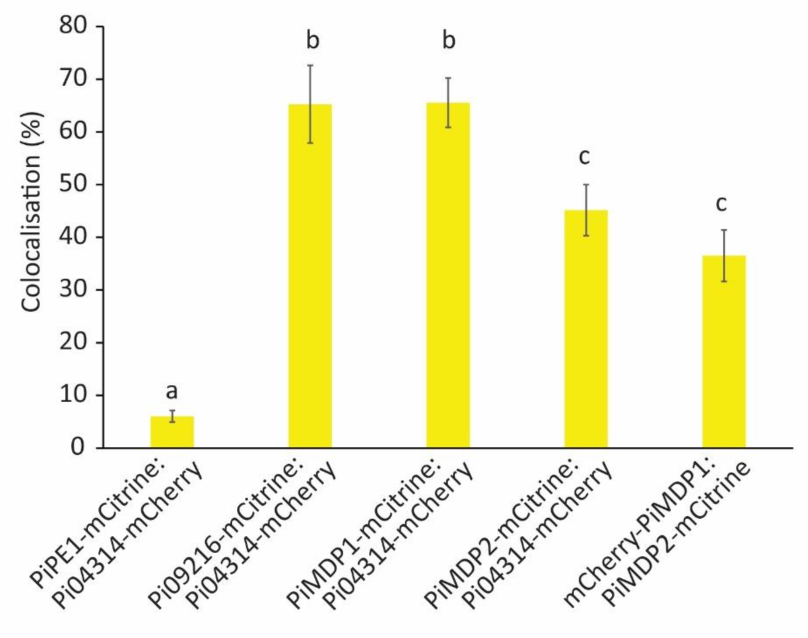


**Figure S9: Colocalisation of MARVEL-domain and RxLR effector proteins in vesicle-like structures in hyphae grown *in vitro*.** Graph shows the percentage of colocalization in *P. infestans in vitro*-grown hyphae for each transformant (as shown in Figure 5). Statistical analysis was carried out using One Way ANOVA with pairwise multiple comparison using the Holm-Sidak method, n ≥12 independent images per transformant from at least two independent experiments. Letters denote significant difference.

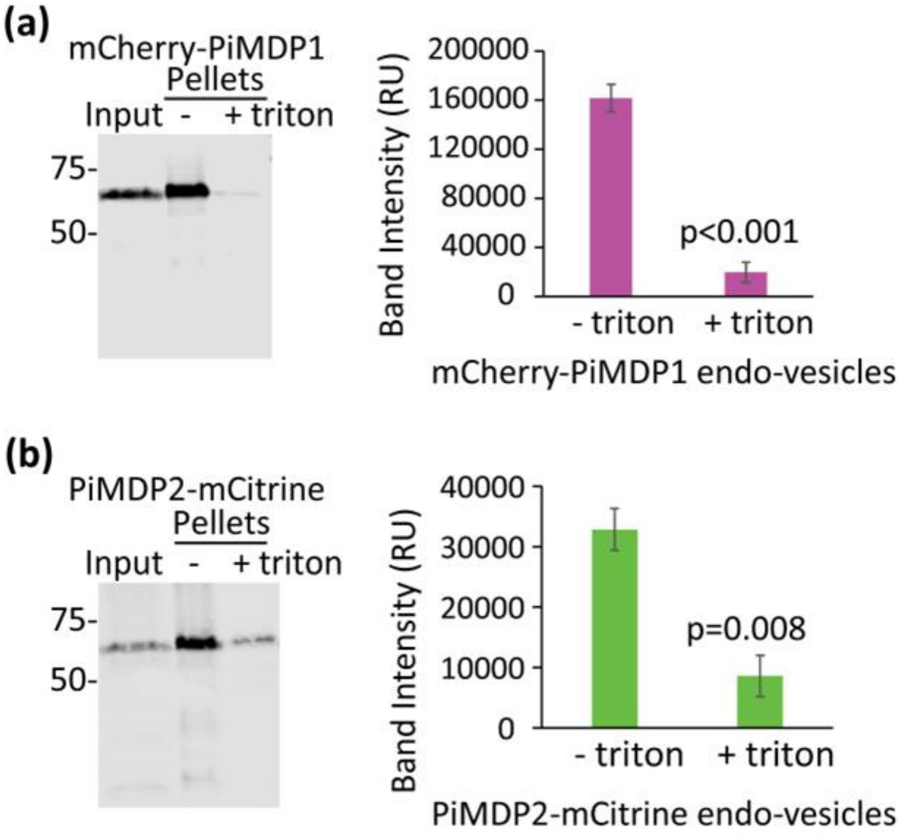


**Figure S10: MARVEL-domain proteins present in endo-vesicle extractions.** Endo-vesicles were extracted from *P. infestans* mycelia expressing (a) mCherry-PiMDP1 and (b) PiMDP2-mCitrine. Western blots show proteins are sensitive to treatment with detergent triton x100, sizes are indicated in kDa. Graphs show quantification of band intensity from three biological replicates plus and minus triton treatments. Error bars are standard error and statistical analysis was done using a Paired T-test on SigmaPlot giving the two-tailed P values shown.

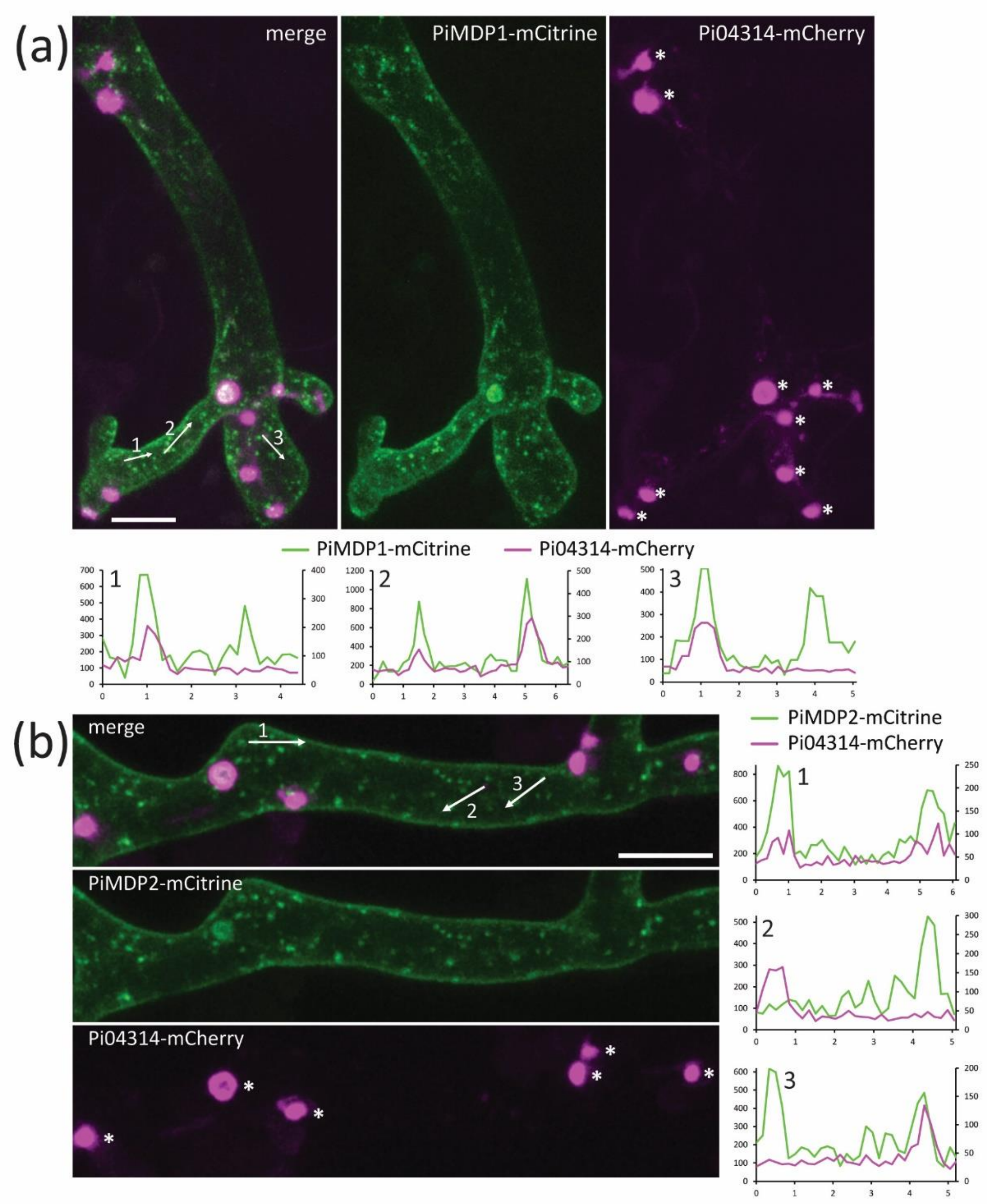


**Figure S11: MARVEL-domain and RxLR effector proteins co-localise in vesicle-like structures in hyphae during infection.**

Further examples of projection images from confocal z series collected from *N. benthamiana* infections by transformants. The mCitrine (green) and mCherry (pink) channels are shown separately and merged with three numbered arrows indicating transects drawn using single optical sections to produce the three numbered fluorescence intensity plots to the right. Where the mCherry intensities were low they are plotted with the right-hand vertical axis to make the patterns more obvious. (a) PiMDP1-mCitrine localises to the plasma membrane (PM) and partially co-localises to vesicles with Pi04314-mCherry. (b) PiMDP2-mCitrine localises to the PM and partially co-localises to vesicles with Pi04314-mCherry. Asterisks indicate haustoria and scale bars are 10 µm. Fluorescence intensity plots show relative fluorescence on the y axis against distance in µm on the x axis.

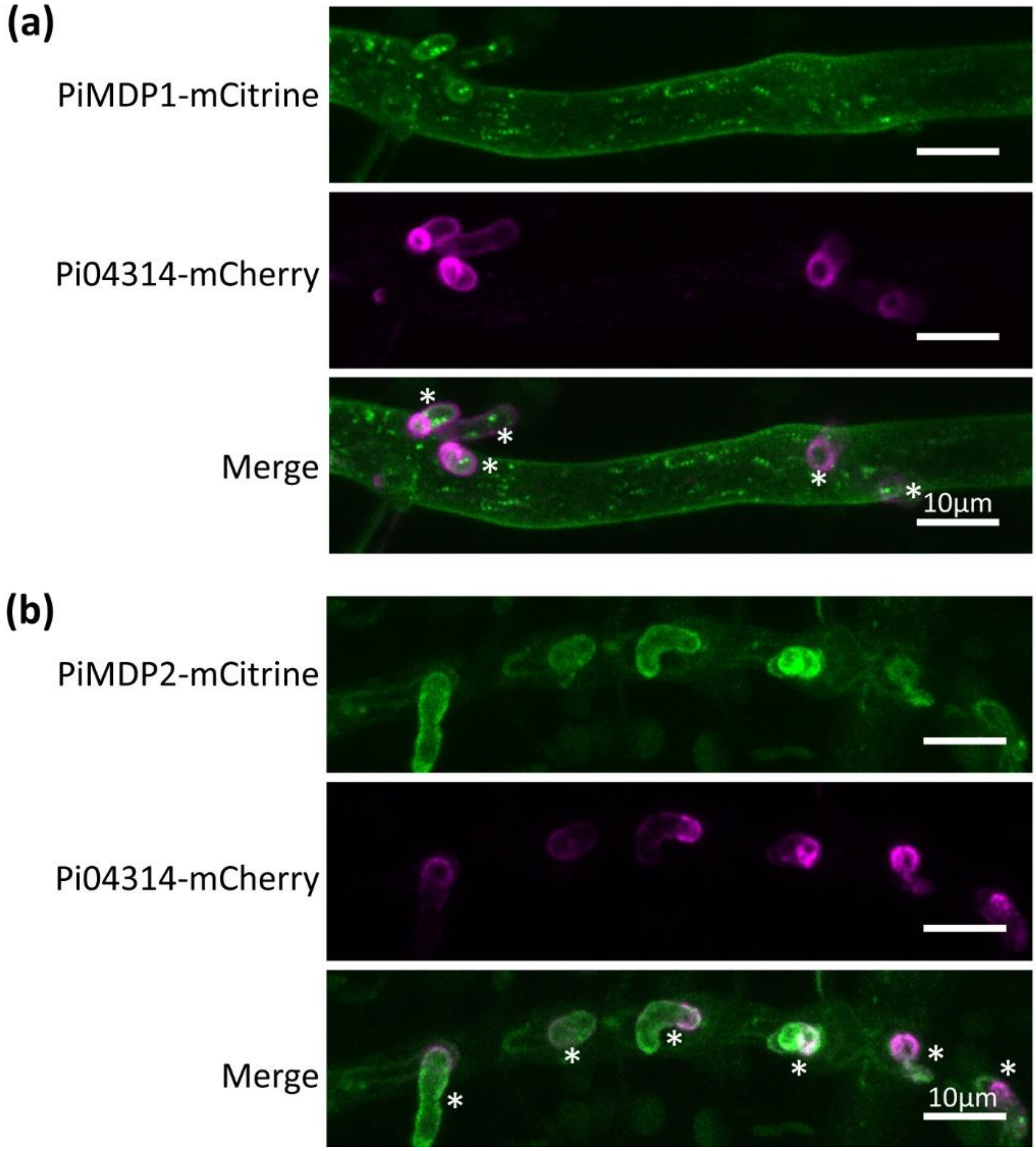


**Figure S12: MDP2 accumulates in the mature haustorial membrane during infection.** Further example images of projections of confocal z series of *P. infestans* hyphae during plant infection showing (a) PiMDP1-mCitrine localised to the plasma membrane (PM) and haustorial membrane (HM) with 04314-mCherry localised around the haustoria and (b) PiMDP2-mCitrine strongly accumulating in the HM with little PM background localisation and 04314-mCherry localised around the haustoria. Asterisks indicate haustoria; the scale bar is 10 μM.
